# Capture-C MPRA: A high-throughput method to simultaneously characterize promoter interactions and regulatory activity

**DOI:** 10.1101/2025.06.11.658967

**Authors:** Coline Arnould, Pia Keukeleire, Fumitaka Inoue, Xiekui Cui, Kelly An, Elizabeth Murray, Xuhuiqun Zhang, Radoje Drmanac, Brock A. Peters, Jay Shendure, Yin Shen, Martin Kircher, Nadav Ahituv

**Author notes:** Correspondence to Martin Kircher and Nadav Ahituv. These authors contributed equally to this work (Coline Arnould, Pia Keukeleire).

## Abstract

*Cis* regulatory elements (CREs) interact with their target promoters over long genomic distances and can be identified using chromatin conformation capture (3C) assays. Their regulatory activity can be functionally characterized in a high-throughput manner using massively parallel reporter assays (MPRAs) that generally test an enhancer alongside a minimal promoter. Here, we developed a novel technology called Capture-C MPRA (ccMPRA) that combines both technologies and can simultaneously obtain chromatin interactions and measure CRE activity alongside their target promoters. We utilized ccMPRA to analyze the regulatory activity of 650 promoters interacting with 42,719 sequences. As C-based techniques also capture isolated promoters, we were able to obtain promoter baseline activity, enabling the identification of both enhancers and silencers. Analysis of CREs interacting with more than one promoter showed significant activity differences depending on the promoter. In summary, ccMPRA can simultaneously characterize chromatin interactions and regulatory activity, allowing to further dissect regulatory grammar.

## Introduction

Mutations in *cis* regulatory elements (CREs) are a major cause of human disease^1–3^. However, our understanding of how these mutations result in specific phenotypes remains limited. There is a wide variety of CREs, including promoters that reside right next to the gene, and distal CREs, such as enhancers (activate gene expression) and silencers (suppress gene expression) that can be located hundreds of kilobases away from the genes they regulate. These CREs act distally by forming three-dimensional (3D) contacts within the nuclear space, establishing chromatin loops that connect them to their target promoter^4^. The formation of these loops is mediated by the coordinated action of CTCF (CCCTC-binding factor) and the cohesin complex^5–8^. Altered chromatin interactions have also been found to be a cause of human disease^9–11^.

CREs are regulated by the binding of various transcription factors (TF) and co-factors and have specific histone modifications based on their activity^12^. Their identification has significantly advanced due to high throughput sequencing-based methods (*e.g.* ChIP-seq, DNase-seq, and ATAC-seq) that have provided genome-wide encyclopedias of millions of regulatory elements in the mammalian genome^13,14^. In addition, Hi-C technology can be used to identify long-range chromatin interactions in a genome-wide manner by capturing closely interacting sequences via ligation^15–18^. Promoter Capture Hi-C (C-HiC) is an adaptation of this technology, utilizing promoter capture sequences to specifically enrich sequences that interact with promoters^19^, providing a more cost-effective way to specifically interrogate promoter-CRE interactions. However, all of these studies are descriptive in nature and do not reveal the function of the promoter interacting sequence.

To functionally characterize regulatory elements *en masse*, massively parallel reporter assays (MPRAs) have been developed. These assays allow the simultaneous testing of hundreds of thousands of sequences and their variants for regulatory activity by placing candidate CREs (cCREs) alongside a transcribed barcode^20^. However, MPRAs have several limitations. These include amongst others the use of short DNA sequences, usually 100-270 base pairs (bp) in length, as cCREs are generally generated using oligosynthesis, which can generate hundreds-of-thousands of short DNA sequences in a cost-efficient manner. Thus, these short-synthesized sequences could miss out on additional longer sequences that are important for regulatory activity. Previous work has shown some MPRA activity differences when testing the same sequence in different lengths^21^. In addition, the majority of MPRAs test for cCRE activity without their target promoter, usually using a minimal promoter, thus not fully capturing the biological activity of the tested regulatory elements. Importantly, MPRAs cannot exhibit the combined regulatory function of CREs with their target promoters and how variants in either one can affect activity and epistatic interactions between them. While some enhancers can activate all promoters equally, it was shown that 10-50% of tested enhancers preferentially activate sub-groups of promoters in mammals, independently of chromatin looping^22–24^. The factors governing this specificity remain unclear, but combinations of transcription factor (TF) motifs may play a role in enhancer-promoter compatibility.

To address these limitations and caveats, we took advantage of the 3D contacts between CREs and promoters to develop Capture-C MPRA (ccMPRA) (**Fig. 1**). This method combines a promoter capture Hi-C followed by the functional characterization of the cCRE directly alongside their cognate promoters via a lentivirus-based MPRA (lentiMPRA)^25^. ccMPRA offers a “two-in-one” approach, simultaneously identifying the target genes of cCREs and assessing their function in combination with their cognate promoter. Using this technology, we captured cCREs interacting with 650 promoters in human hepatocytes (HepG2) via Capture Hi-C and assessed their regulatory activity in combination with each of their interacting regions by directly cloning the capture Hi-C library into a lentiMPRA library. An artifact of capture Hi-C is that promoter fragments get captured by themselves^26^. However, by retaining these sequences in our ccMPRA analyses, we were able to establish baseline promoter activities thus allowing us to identify not only enhancers but also silencers of that promoter activity. In total, we identified 886 enhancers and 1,251 silencers, including 1,573 that had not been previously annotated for this cell line in ENCODE. We also identified numerous sequences that interacted with more than one promoter, some having differential activity (including both enhancing and silencing) depending on the promoter, suggesting that regulatory activity differs based on the interacting promoter. Additionally, we measured the regulatory activity of the short length proportion of cCREs, i.e. those between 200 and 270 bp long, from our ccMPRA in an oligo-synthesized lentiMPRA that contains a minimal promoter, finding an overall good correlation between both technologies and further analyzed sequences that showed difference between them. In summary, we developed a novel MPRA that allows capturing CREs alongside their target promoters, identifies not only enhancers but also a larger proportion of silencers, and furthermore shows that the identity of the target promoter can have a significant effect on CRE activity.

**Fig. 1:**
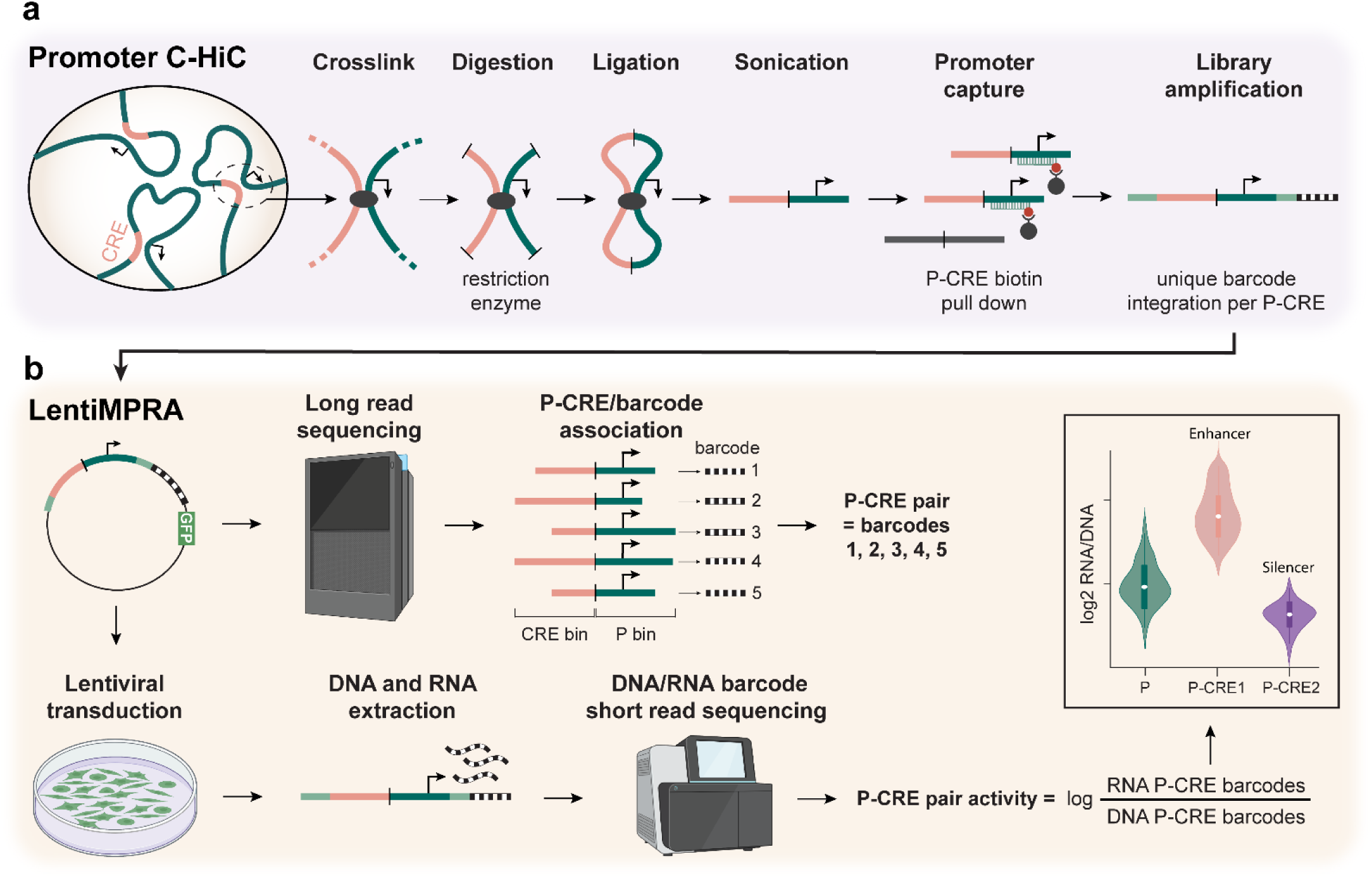
Capture-C MPRA. **a**, Capture Hi-C (C-HiC) identifies significant 3D interactions between promoters (P) and distal *cis*-regulatory elements (CREs). DNA interactions are crosslinked, followed by DNA digestion with a restriction enzyme. A ligation step then generates circular DNA hybrids containing the two interacting DNA regions. Following sonication, the linearized DNA undergoes a promoter capture step, where biotinylated RNA probes hybridize to promoters, enabling a biotin-streptavidin pull-down of P-CRE pairs. P-CRE pairs are then amplified and unique 15 base pair barcodes are added to each pair through PCR. **b**, The promoter C-HiC library (P-CRE pairs with barcodes) is cloned into a lentivirus-based massively parallel reporter assays (lentiMPRA) vector in front of a green fluorescent protein (GFP) to enable the assessment of regulatory activity of thousands of DNA sequences within their chromatin context. Long-read sequencing is performed to link each P-CRE sequence to their unique barcodes. As sequences differ in length for each P-CRE interaction, the sequences are binned to provide numerous barcodes per interaction. The library is transduced into cells via lentivirus and after three days, DNA and RNA are extracted, and DNA/RNA barcodes are sequenced using short-read sequencing. Regulatory activity is quantified as the log ratio of RNA to DNA barcodes for each P-CRE pair and compared to signals from the corresponding promoters alone (in several instances promoter fragments do not ligate to an interacting region, a common C-HiC artifact). Some elements in the figure were created with BioRender.com.

## Results

### Development of a customized capture Hi-C protocol

To capture CREs with their target promoter, we first developed a customized promoter Capture Hi-C (C-HiC) protocol that enriches for promoters and captures longer promoter-interacting sequences. We used the Arima Capture-HiC+ kit with a custom probe panel (Arima Genomics) to enrich chromatin interactions for 650 selected promoters and their interacting genomic regions (**Fig. 1a**). This set included 100 positive controls (highly interactive promoters with strong transcriptional activity in HepG2 cells), 100 negative controls (weakly interactive promoters with low transcriptional activity in HepG2 cells), and 450 test promoters (promoters selected among the most interactive in HepG2 cells). Since promoter lengths can vary from 100 to 1000 bp and CRE lengths can range from 10 to 1000 bp^27,28^, we adjusted the C-Hi-C protocol to create larger DNA fragments, thereby increasing the likelihood of obtaining complete promoter and cCRE sequences in the final library. This adjustment involved using a single restriction enzyme instead of the recommended two (reducing DNA digestion) along with gentler sonication (see Methods for more details). As a result, the Hi-C pairs ranged from 500 bp to 2.5 kb, with an average peak size of 1,300 bp instead of the recommended 400 bp (**Extended Data Fig. 1a**). To enhance the capture and pull-down of these large Hi-C pairs, we initiated the capture using a larger input of DNA from the Hi-C library. In the final step, a unique 15 bp random barcode was introduced to each Hi-C pair via PCR using custom primers. These primers also included sequences homologous to the MPRA plasmid, which allow subsequent cloning by recombination.

The C-HiC library, containing captured promoters ligated to their interacting regions (i.e. cCREs) and their barcodes, were cloned into the lentiMPRA vector upstream to an EGFP reporter gene (**Fig. 1b**). Unlike standard MPRA, this construct does not contain a minimal promoter. Notably, each promoter and cCRE within the Hi-C pair can be inserted in forward or reverse orientation and at either position, with the promoter placed upstream or downstream of the cCRE. Unless specified otherwise, the MPRA analysis accounted for the promoter’s orientation (forward or reverse) in each pair, considering only the properly oriented promoters (forward insertion for sense promoters and reverse insertion for antisense promoters), while considering the orientation of the cCRE and the relative positioning of the promoter and cCRE (upstream or downstream) indistinctly. To associate each promoter-CRE (P-cCRE) pair with its corresponding barcode, the promoter-cCRE-barcode part of the constructs were amplified by PCR and sequenced using long-read sequencing (PacBio Revio and Complete Genomics; see Methods). As the lengths of the sequences vary between different P-cCRE pairs, sequences were binned into loci, i.e. sequences having the same promoter and overlapping cCRE were considered to be from the same locus and used to increase the number of observations for the specific interaction, i.e. providing multiple barcodes in analysis.

Since multiple steps of the promoter C-HiC protocol were customized to accommodate the ccMPRA application, we first sought to evaluate capture performance. Notably, the length of C-HiC pairs was significantly greater than that of regular C-HiC (median = 710 bp, mean = 741.75 bp instead of ∼300 bp for a regular C-HiC; **Extended Data Fig. 1b**). This increased length reflects the combined contributions of promoter and cCRE sequences within the C-HiC pairs, with promoters ranging between 30 and 3023 bp and having a median length of 553 bp (mean = 588.19 bp) and cCREs ranging between 30 and 3089 bp and with a median length of 254 bp (mean = 330.51 bp; **Extended Data Fig. 1c**). Despite the protocol modifications, initial quality control demonstrated a high percentage of valid Hi-C pairs (98.68%, **Extended Data Fig. 1d**) and strong capture efficiency, with 79% of valid Hi-C pairs containing at least one targeted promoter (**Fig. 2a**, **Extended Data Fig. 1e**). As expected, positive control promoters exhibited a higher number of interactions compared to negative control promoters (**Extended Data Fig. 1e-f**), further validating both the specificity of our approach and the design of promoter categories.

**Fig. 2:**
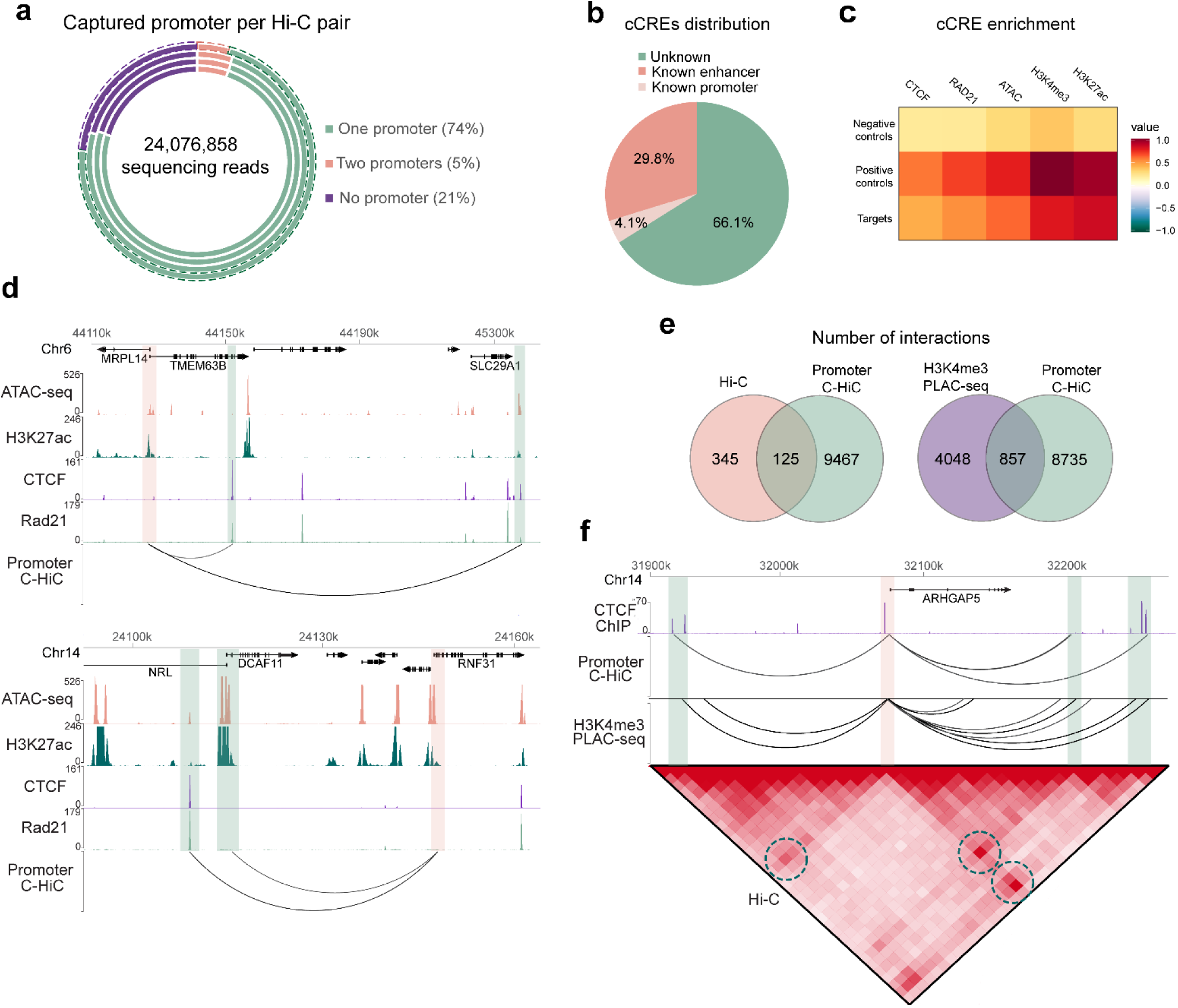
Custom promoter C-HiC validation. **a**, On-target rate of the promoter C-HiC, i.e. the proportion of C-HiC pairs containing successfully captured promoters (either one target promoter or 2 interacting target promoters), and the proportion of C-HiC containing no target promoter. The circles represent the results obtained from four long read sequencing runs (3 Pacbio Revio and 1 Complete Genomics (outer dashed circle)) with the average percentage of all four sequencing results indicated for each category. **b,** Percentage of the promoter interacting regions, considered as candidate *cis* regulatory elements (cCREs) overlapping with ENCODE SCREEN^14^ v3 annotated CREs. **c,** Enrichment of ChIP-seq for CTCF, RAD21, H34Kme3, H3K27ac and ATAC-seq signals for negative controls, positive controls and sequences captured by the target promoters (“Targets”). **d,** WashU Epigenome Browser^34^ screenshots showing HepG2 ATAC-seq, H3K27ac/CTCF/RAD21 ChIP-seq signals and the promoter C-HiC signal (arcs representing interaction) for the *TMEM63B* and *RNF31* loci. The orange shaded region indicates the target promoter and green depict the regions interacting with this promoter. **e,** Overlap of chromatin interactions between the promoter C-HiC library and previously published Hi-C^14,30–32^ and H3K4me3 PLAC-seq^33^. **f,** WashU Epigenome Browser^34^ screenshot comparing interactions identified in the promoter C-HiC and HepG2 H3K4me3 PLAC-seq^33^ and Hi-C^14,30–32^, as well a CTCF ChIP-seq signal for the ARHGAP5 locus. Circles indicate matching interactions in all three datasets.

We used the CHiCAGO pipeline^29^ to call significant interactions in our C-HiC and found 9,592 significant interactions (score>5, 5kb resolution; interactions ≤2Mb). To confirm the ability of our custom promoter C-HiC protocol to capture putative enhancers, we assessed the overlap between the promoter interacting regions (cCREs) and ENCODE-annotated enhancers, finding an overlap of 29.8% (**Fig. 2b**). Supporting an efficient capture of distal regulatory elements, we observed a strong enrichment of cCREs interacting with positive control promoters and a very low enrichment of cCREs interacting with negative control promoters for enhancer-associated chromatin features (**Fig. 2c-d**). These include HepG2 ChIP-seq for CTCF and the cohesin subunit Rad21, histone marks such as H3K27ac and H3K4me3 and ATAC-seq (**Fig. 2c-d**). To further validate the performance of our customized C-HiC protocol, we compared the interactions identified by our method with those detected using established approaches, including Hi-C^14,30–32^ and H3K4me3 PLAC-seq^33^ from HepG2 cells. Despite differences in experimental protocols and computational pipelines for interaction calling, we observed a substantial overlap between the datasets, 26% of the Hi-C interactions and 17.5% of the H3K4me3 PLAC-seq interactions involving a promoter from our set of captured promoters overlapped our C-HiC (**Fig. 2e-f**), supporting the robustness of our capture approach. Altogether, our data confirms that our customized promoter C-HiC protocol captures valid chromatin interactions.

### ccMPRA identifies functional regulatory sequences

Following validation of our C-HiC library, we carried out lentiMPRA to assess the regulatory activity of these Hi-C pairs. The library was packaged into lentivirus and transduced into HepG2 cells to provide an ‘in genome’ readout which was shown to be more strongly correlated with sequence-based models and ENCODE annotations and to provide higher predictions of cell-type specificity compared to episomal MPRA^35,36^. Transduction efficiency was assessed by GFP expression in cells and the multiplicity of infection (MOI) was measured by qPCR (MOI ∼100; see Methods). The transcriptional activity of each promoter-CRE pair was quantified by sequencing RNA and DNA barcodes with two biological replicates containing three technical replicates each. RNA barcode counts were normalized against DNA barcode counts, accounting for barcode differences in construct representation and integration.

We first examined the number of barcodes per C-HiC pair. We obtained a median of 4.4 barcodes per identical sequence (i.e. sequences that had the exact same promoter and/or captured sequence) for the first biological replicate and 4.7 for the second biological replicate (**Extended Data Fig. 3a**). This corresponds respectively to an average recovery of 61.8% and 57.7% of the barcodes identified through the promoter-CRE barcode association long-read sequencing. To assess technical reproducibility, we examined the correlation between replicates within each biological replicate. Specifically, intra-assay Pearson correlations for pairwise comparisons of activity scores (logRNA/DNA barcodes per C-HiC sequence) exceeded 0.63 across all assays (**Extended Data Fig. 2a-b**), despite the high complexity of our library and limited number of barcodes per sequence.

Given the variability in C-HiC pair sequence length (**Extended Data Fig. 1b-c**), we binned regions by restriction fragments before MPRA analysis, focusing only on paired-bins containing properly oriented promoters (**Extended Data Fig. 3b**). To ensure reliability, we retained only paired-bins with ≥ 5 barcodes in the MPRA, whether due to multiple sequences or to multiple barcodes per sequence, resulting in a median of 8 and mean of 12.75 barcodes per bin and a median of 2 and average of 2.15 sequences per bin (**Extended Data Fig. 3c-d**). Filtered paired-bins had a median bin size of 545 bp and a mean size of 557.4 bp (**Extended Data Fig. 3e**). After categorizing bins into promoter and cCRE bins, we obtained a promoter median size of 446.5 bp, mean of 494.6 bp and maximum of 1,614 bp, and a cCRE median bin size of 133 bp, mean of 149.8 bp and maximum of 1,654 bp (**Extended Data Fig. 3f**). Positive control promoter bins were found to capture more interacting regions/cCREs than negative controls (**Extended Data Fig. 3g**), further confirming the specificity of the protocol.

We next assessed the efficiency of the lentiMPRA. Categorizing the paired-bins based on the presence or absence of a transcription start site (TSS) found that paired-bins containing a TSS exhibited stronger MPRA activity (**Extended Data Fig. 4a**). Next, we assessed the range of activity scores across different types of paired-bins, including positive and negative promoter controls, as well as target promoters interacting with other promoters or with non-promoter regions (“other”), corresponding to the cCREs of interest. Paired-bins containing a positive control promoter exhibited higher MPRA activity than those with a negative control promoter, further validating the efficiency of the lentiMPRA (**Fig. 3a**).

**Fig. 3:**
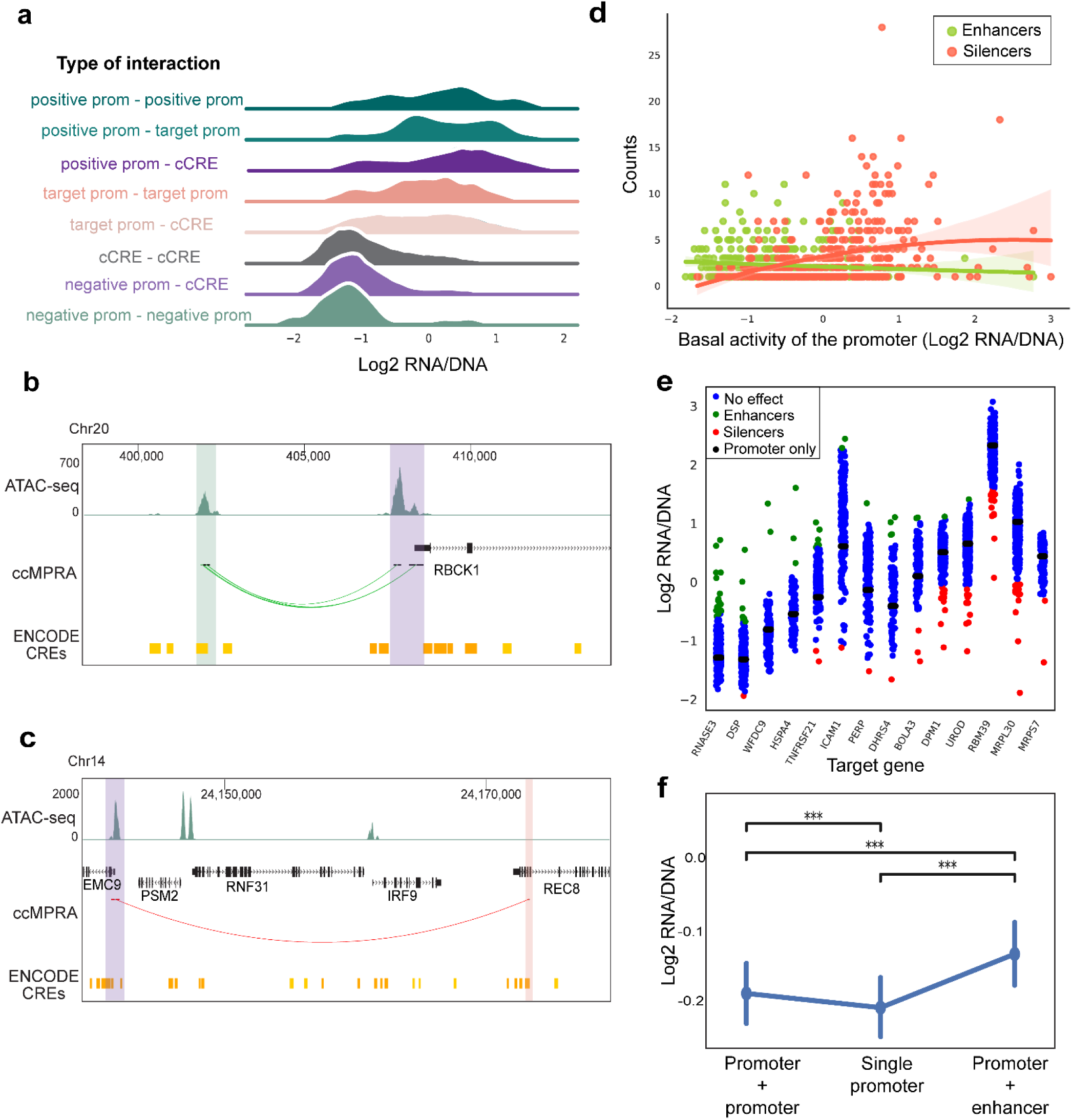
lentiMPRA regulatory activity of C-HiC sequences. >**a**, The distribution of log2(RNA/DNA) ratios for positive and negative control promoters (prom), as well as the target promoters, categorized by their interactions with other promoters or candidate *cis* regulatory elements (cCREs). **b-c,** UCSC Genome Browser screenshot showing an example of an enhancer identified by ccMPRA for RBCK1 (**b**) or a silencer for EMC9 (**c**). HepG2 ATAC-seq and CREs from ENCODE^31^ are shown alongside our ccMPRA signal (green and red arcs represent an enhancing or silencing interaction respectively). Purple shade indicates the target promoter and the green/red shade depicts the enhancer or silencer region. **d,** Plot showing the distribution of the number of enhancers (green dots) or silencers (red dots) identified per promoter as a function of the promoter’s basal activity. **e,** Examples of cCREs regulatory activity distribution as compared to the basal activity of the promoter alone (black dot) for a subset of promoters. Blue dots represent cCREs without significant activity effects, green and red represent significant enhancers or silencer activities respectively (z-score >2 or <-2 respectively; see Methods). **f,** Log2(RNA/DNA) representing the MPRA activity of single promoters versus those same promoters interacting with another promoter or with enhancers (median activity of all the interacting enhancers) (n=843 promoter bins analyzed). The vertical lines represent the 95% confidence interval and *** represents p<0.0001 using a paired t-test.

### ccMPRA identifies both enhancers and silencers

A well-known artifact of C-based techniques is the capture of isolated promoter fragments that did not ligate to other sequences^26^. We leveraged this phenomenon to compare the activity of P-cCRE pairs with that of the promoter alone, enabling us to infer cCRE activity (z-score; see Methods). This approach allowed us to identify not only enhancers but also silencers. We classified cCRE bins as significant enhancers or silencers if their z-score was >2 or <-2 respectively. Using this approach, we identified 886 enhancers and 1,251 silencers, underscoring the advantage of using promoters with varying basal activities to uncover both activating and repressive CREs, in contrast to using a minimal promoter alone (**Fig. 3b-c**, **Extended Data Fig. 4b-c**). Interestingly, our data suggest that highly expressed promoters primarily interact with silencers, whereas lowly expressed promoters mainly interact with enhancers (**Fig. 3d**, **Extended Data Fig. 4d**, **Supplementary Fig. 1**). To further illustrate this, we provide an analysis of a subset of promoters with variable self-activity (black dots) alongside the distribution of their associated cCRE regulatory activity, showing an increase in enhancers for weak promoters and strong promoters having more silencers (**Fig. 3e**). Of note, this observation could also be due to a limitation of our technique, potentially failing to detect activity decreases when promoter activity is already very low, and conversely, increases when activity is high. However, the relationship between promoter intrinsic activity and the number of detected enhancers or silencers is not strictly linear, as evidenced by the scatter distribution and trend lines, indicating that this pattern cannot be attributed solely to technical limitations (**Fig. 3d**). Nevertheless, the observed pattern, whereby stronger promoters exhibit reduced responsiveness to further activation and weaker promoters show heightened responsiveness, is consistent with previously reported studies^22–24,37^.

We also observed that 60 non-targeted promoters acted as enhancers, in line with previous studies showing promoters can function as enhancers of other promoters^38–41^. Specifically, when we measured the activity of promoters alone or in interaction with another promoter, we observed that the interaction with another promoter enhanced transcriptional activity, though to a lesser extent than enhancers (n= 843 promoter bins, **Fig. 3f**).

### Architectural determinants of CRE activity

We then examined the architecture of distal CRE–promoter communication. As mentioned above, our assay can measure cCRE activity when placed either upstream or downstream of its cognate promoter. We observed a very high correlation between upstream and downstream configurations, indicating that many CREs regulate transcription equally well on either side of the promoter (**Extended Data Fig. 5a**). This is in line with previous reports that enhancers are largely position-independent as long as they can physically contact their target promoter by chromatin looping^42^.

Next, we assessed the impact of orientation. To simplify comparisons, we initially fixed each promoter in its endogenous orientation. We then evaluated the activity of each promoter and cCRE in both forward and reverse orientations. Consistent with prior studies, distal elements, unlike promoters, generally function in an orientation-independent manner (**Extended Data Fig. 5b**)^22,43–45^. However, our data suggested that promoters interacting with silencers exhibited greater dependence on orientation than promoters interacting with enhancers (**Extended Data Fig. 5c-d**). A possible explanation is that repressive complexes or orientation-sensitive transcription factor binding sites at the promoter may require a specific orientation to be regulated by silencers effectively, whereas enhancer-bound activators endure bidirectional contact. We then examined whether the number of preexisting promoter-cCRE contacts (as measured by promoter C-HiC) predicted CRE activity, observing no significant correlation (**Extended Data Fig. 5e**). This likely reflects that Hi-C contact frequency reports structural proximity rather than intrinsic regulatory potential, so regions with many contacts are not necessarily more enriched for functional enhancers or silencers.

Finally, we assessed whether CRE length or endogenous genomic distance to the target promoter influenced regulatory activity. Interestingly, we found that silencers tend to lie closer to their target promoters than enhancers (**Extended Data Fig. 5f**). However, we found no significant correlation between CRE length or endogenous genomic distance to the target promoter with regulatory activity (**Extended Data Fig. 5g-h**). Recent work has shown that long-range enhancer function is determined predominantly by intrinsic regulatory strength^24^, implying that genomic distance has minimal impact on activity and that chromatin looping readily bridges large intervening gaps to bring enhancers into proximity with their target promoter^4,46^.

### Regulatory features of enhancers and silencers

We next characterized the regulatory features of enhancers and silencers. We first characterized the distribution of various histone modifications using HepG2 ChIP-seq data from ReMap 2022^47^. After excluding the annotated promoters that were detected as enhancers or silencers, we observed an enrichment for enhancers with a variety of HepG2 active histone marks (K3K27ac, H3K9ac, H2AFZ) which had lower enrichment levels in silencers (**Fig. 4a**, **Extended Data Fig. 6a**). In contrast, silencers had stronger enrichment for known silencer marks such as H3K79me2 and H4K20me1 (**Fig. 4a**, **Extended Data Fig. 6a**). The overlap between silencers and the repressive marks H3K79me2 and H4K20me1 is in line with a previous report that carried out high-throughput silencer assays and found enrichment for these marks along with H3K36me3^48^, which we also observed in our silencers. Additionally, we found that both enhancers and silencers are enriched for H3K4me3, a histone modification typically associated with active promoters or enhancers^49^. This unexpected enrichment on silencers likely reflects the subset of silencers located near promoters (median distance between promoter and H3K4me3-overlapping silencers = 2,769 bp; non-overlapping = 70,760.5 bp), where promoter-associated chromatin signatures may extend into nearby regulatory elements. Surprisingly, we also detected an enrichment of the enhancer-specific histone modifications H3K27ac and H3K9ac on silencers as well, albeit to a lesser extent than enhancers. While unexpected, this result is supported by recent studies showing that certain silencers can carry histone modifications typically associated with active enhancers^48,50–53^. It is indeed possible that silencers reside at the vicinity of activating sequences^54^, reflecting the existence of global regulatory regions, in which distinct sub-elements exert either activating or repressive effects on transcription, resulting in a fine-tuned control of gene expression. Finally, both enhancers and silencers were equally enriched for H3K36me3, supporting the established dual role of this histone mark^55^. We also found enhancers to be more enriched for open chromatin (DNase hypersensitive sites and ATAC-seq in HepG2 cells) than silencers (**Fig. 4a**, **Extended Data Fig. 6a**). This limited overlap of silencers with accessible chromatin is in agreement with what was observed in *Drosophila* using Silencer-seq, an unbiased method to detect silencers^43^.

**Fig. 4:**
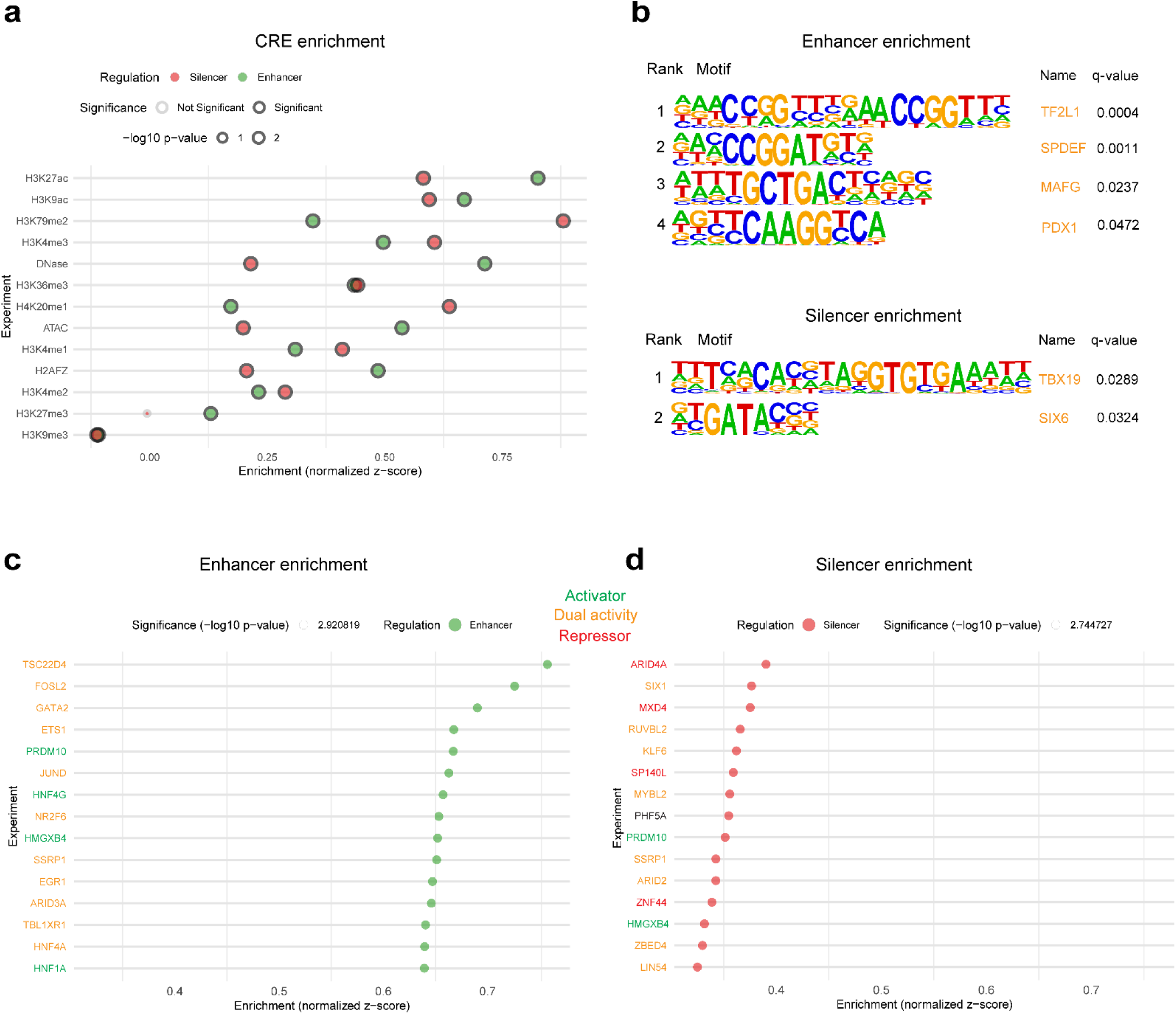
Features of ccMPRA CREs. >**a**, ATAC-seq, DNAse hypersensitive sites and ChIP-seq (histone modifications) signal enrichment from ENCODE (HepG2 cells) is shown across enhancers and silencers identified by ccMPRA. Circle size represents -log10 p-value of permutation tests. **b,** Transcription factor (TF) motifs significantly enriched for enhancer or silencer sequences, detected using Homer^56^ along with their name and q-value. **c-d,** Enrichment of ChIP-seq signal analyzed from ReMap 2022^47^ (HepG2 cells) for various transcription factors for enhancers (**c**) or silencers (**d**). The top 15 enriched TF are shown and green, yellow and red TF names represent TFs that are known to be activators, have dual activity or repressor activity respectively. Z-scores represent the distance between the evaluation of our CRE regions and the mean of the random evaluations divided by the standard deviation of the random evaluations of the permutation test.

To investigate which TFs may regulate our CREs, we carried out TF motif enrichment using HOMER^56^. We found that enhancer-associated CREs were significantly enriched for motifs recognized by the dual-activity factors TF2L1, SPDEF, MAFG and PDX1 (**Fig. 4b**)^57–62^. In contrast, silencer-associated CREs were enriched for motifs corresponding to TBX19 and SIX6 (**Fig. 4b**), both of which have been reported to exhibit context-dependent activating and repressive functions^63,64^.

As ReMap 2022^47^ integrated ChIP-seq datasets for the binding of hundreds of TFs in HepG2 cells, we used these datasets to determine the top TFs that preferentially bind to enhancers or silencers. We found that enhancers are mainly bound by TFs that were characterized as transcriptional activators (PRDM10, HNF4G, HMGXB4, HNF1A)^65–68^, as well as TFs known to have a dual activity depending on context (TSC22D4, FOSL2, GATA2, ETS1, JUND, NR2F6, SSRP1, EGR1, ARID3A, TBL1XR1, HNF4A) (**Fig. 4c**)^66,69–80^. Several of the transcription factors identified are canonical liver-enriched regulators (HNF1A, HNF4A, HNF4G and GATA2), which is consistent with the hepatic origin of the HepG2 cell line used in this study. Conversely, silencers were enriched for TFs with established repressor activity (ARID4A, MXD4, SP140L, ZNF44)^81–83^ or activator activity (PRDM10, HMGXB4)^65,67^ and for TFs having context dependent dual activity (SIX1, RUVBL2, KLF6, MYBL2, SSRP1, ARID2, ZBED4, LIN54)^76,77,84–93^ (**Fig. 4d**). Altogether, our data demonstrate that enhancers and silencers exhibit both distinct and shared features.

Of the regulatory elements we identified, 73.6% (614 enhancers and 959 silencers) were not previously annotated as cCREs by ENCODE SCREEN^31^ v3, which primarily relies on chromatin accessibility and histone modifications to determine regulatory potential (**Extended Data Fig. 6b**). Since SCREEN does not distinguish silencers from enhancers, we next examined the subset of enhancers identified by our assay. We observed that these sequences have significant enrichment for canonical enhancer-associated histone modifications (H3K27ac, H3K9ac, and H3K4me1), but to a lesser extent than ENCODE-annotated enhancers (**Extended Data Fig. 6c**), suggesting that ccMPRA might be more sensitive in its ability to detect regulatory elements with weaker chromatin signatures. Of note, we observed no substantial difference in regulatory activity between these two categories (average z-score: 2.61 for ENCODE enhancers vs. 2.45 for non-ENCODE enhancers), suggesting that the non-ENCODE enhancers are unlikely to be false positives.

### Promoters affect CRE activity

Several of our CREs were captured by multiple promoters. We thus set out to test whether different promoters can have an effect on CRE activity. For CREs ligated to more than one promoter that passed our ccMPRA results filters, we had 2,093 that interacted with promoter bins of at least two genes, 78 with promoter bins of at least three genes and 5 with promoter bins of at least four genes (**Fig. 5a**, **Extended Data Fig. 7a**). We observed significant activity differences depending on the target promoter with differences in normalized activity scores ranging between 0.001 and 6.73. In addition, we also saw CREs that function as enhancers with one promoter and silencers with another (**Fig. 5a**, **Extended Data Fig. 7a**). Analyses of these multi-promoter interacting CREs found them to be enriched for canonical enhancer or silencer-associated histone marks (H3K27ac, H3K9ac, H3K4me3 and H3K79me2, H4K20me1 respectively) and to be more accessible than CREs interacting with a single promoter (**Fig. 5b**). We hypothesized that physically bridging two or more promoters requires more robust ‘scaffolding’ than a single chromatin loop. Indeed, we found that these multi promoter interacting CREs were also enriched for known regulators of chromatin looping, such as CTCF and members of the cohesin complex (**Extended Data Fig. 7b**). Overall, our data indicate that CREs interacting with multiple promoters act as dynamic regulatory hubs whose activity is fine-tuned by promoter context which depends on an enhanced complement of both activating/repressive chromatin features and architectural proteins to coordinate multi-gene expression controls.

**Fig. 5:**
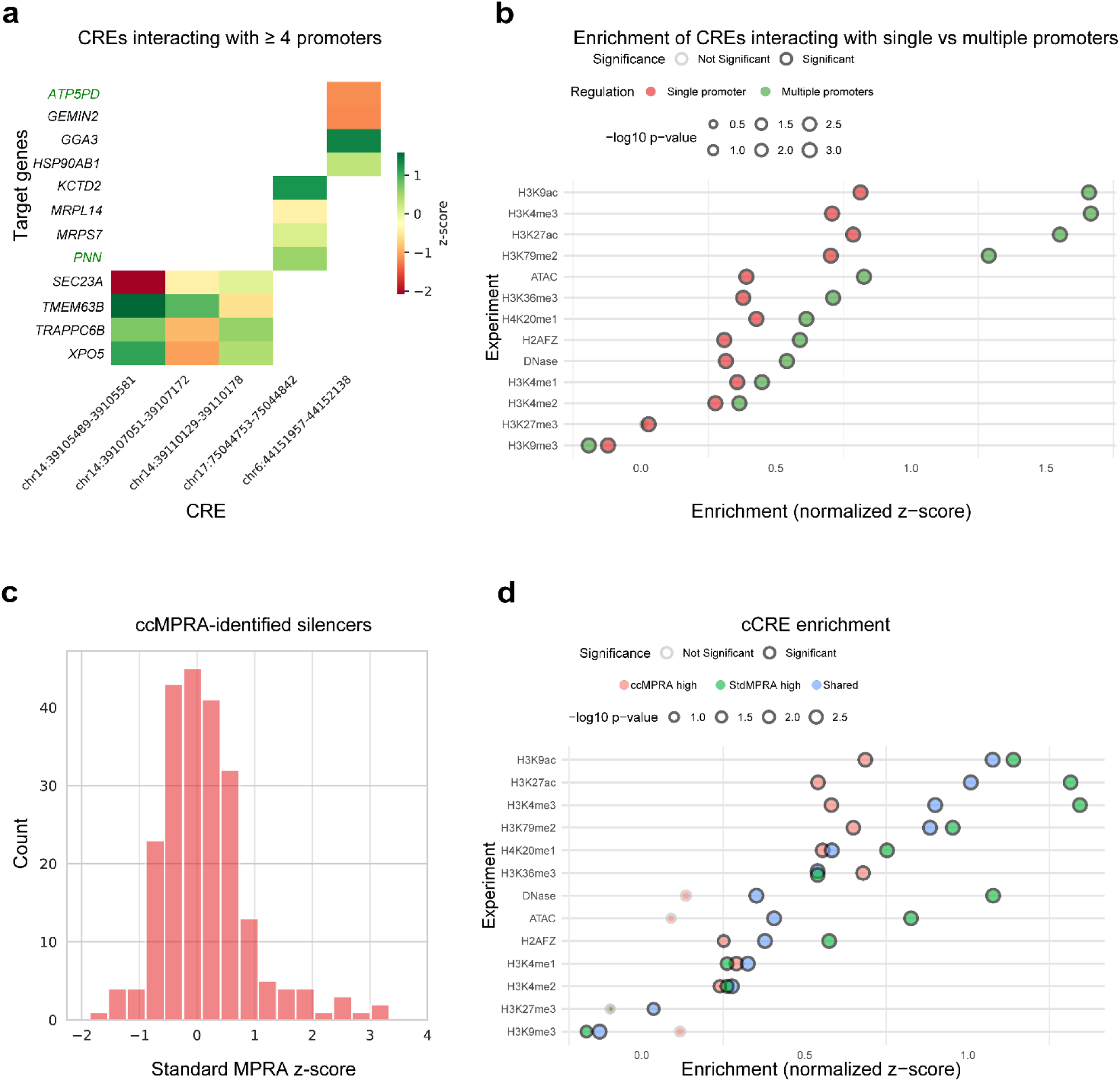
Promoters affect CRE activity. >**a**, Heatmap showing the regulatory activity (z-score) of CREs interacting with four or more promoters, plotted separately for each promoter they contact. **b,** ATAC-seq, DNAse hypersensitive sites and ChIP-seq (histone modifications) signal enrichment from ReMap 2022^47^ (HepG2 cells) is shown across CREs identified by ccMPRA, whether they interact with a single promoter (red) or with multiple promoters (green). Circle size represents -log10 p-value of permutation tests. **c,** Distribution of the number of silencers identified by ccMPRA as a function of their z-scores obtained in the standard MPRA assay. **d,** ATAC-seq, DNAse hypersensitive sites and ChIP-seq (histone modifications) signal enrichment from ReMap 2022 (HepG2 cells) is shown across cCREs, whether they have a higher detected activity in the ccMPRA (ccMPRA high, red), in the standard MPRA (StdMPRA high, green) or have a similar activity between the two assays (Shared, blue). Circle size represents -log10 p-value of permutation tests.

### Comparison of ccMPRA to a standard minimal promoter MPRA

To characterize the differences between ccMPRA and standard MPRAs that have a minimal promoter, we generated a standard MPRA library with all ccMPRA loci that were 200-270 bp in length (6,231 sequences). These were cloned to a lentiMPRA vector containing a minimal promoter and then used a standard lentiMPRA protocol (see Methods)^25^ to test their activity in HepG2 cells. To control the experiment, we added 100 positive controls and 100 negative controls that were shown to have high or low activities respectively in a previous MPRA in HepG2 cells^94^, and added 200 shuffled sequences to estimate the basal activity of the MPRA. We obtained a median of 1,372 barcodes per sequence and obtained sufficient data, after various data quality filters (see Methods), for 5,963 sequences that showed strong correlations between the three technical replicates (average Pearson’s r=0.96 between replicates) (**Extended Data Fig. 8a-b**). As expected, the positive controls exhibited a higher regulatory activity than the negative controls and shuffled sequences (**Extended Data Fig. 8c**). We then compared results between the standard lentiMPRA and the ccMPRA for the same sequences. Overall, we found a good correlation between experiments (**Extended Data Fig. 8d-e**). We also observed sequences that showed higher activity in either assay. We first focused on the sequences that displayed higher regulatory activity in the standard MPRA compared to the ccMPRA (“stdMPRA high”, n=168). Interestingly, these stdMPRA-high sequences had an average z-score of –2.09 in the ccMPRA, indicating that many of them could function as silencers that are not detected by the standard MPRA. Supporting this idea, we examined the distribution of activity scores in the standard MPRA for the silencers identified by ccMPRA and found that the majority exhibited little to no regulatory effect, while a small subset showed enhancer activity (**Fig. 5c**). These results further highlight one of the key advantages of ccMPRA, the ability to functionally characterize both enhancers and silencers in a promoter-dependent context.

Next, we analyzed the sequences that displayed higher regulatory activity in the ccMPRA compared to the standard MPRA (“ccMPRA high”, n=210). These sequences showed enrichment for a variety of transcription factors, to a same or slightly lower extent than sequences that have similar regulatory activity between the two assays (“shared”, n=5,585) (**Extended Data Fig. 8f**). Similarly, these “ccMPRA high” sequences also showed enrichment for a variety of histone marks associated with regulatory activity that was lower than that of “shared” and “stdMPRA high” (**Fig. 5d**). Together, these findings suggest that ccMPRA might be more sensitive for sequences with reduced regulatory marks, as previously observed for non-ENCODE annotated sequences. This observation could also be due to these cCREs being tested with their target promoter instead of a minimal promoter. It is important to also note that these findings could also be related to a variety of technical differences between these assays.

## Discussion

To overcome two main caveats in MPRAs, the use of short sequences and a minimal promoter, we developed a novel MPRA that utilizes promoter capture Hi-C to clone longer sequences along with their target promoters into MPRA vectors. By customizing the promoter capture Hi-C, we were able to enrich for promoter-containing sequences and longer lengths. Due to the inherent capture of promoter fragments by themselves via promoter capture Hi-C^26^, we could quantify basal promoter activity and were able to infer not only enhancing effects but also a substantial proportion of silencers via ccMPRA. Analysis of these sequences found enhancers to have an enrichment for TFs and histone marks fitting with an active role versus silencers that were enriched for histone marks and TFs involved with more of a repressive role. As our promoter capture Hi-C also captured CREs with multiple different promoters, we were able to observe differential CRE activity depending on the captured promoter. Comparison to a minimal promoter based MPRA with CREs similar in size found overall similarities between the results with differences for silencers and ccMPRA identifying active sequences that have lower regulatory mark enrichment.

Our modified promoter Capture Hi-C protocol allowed the cloning of longer sequences and enriched for more sequences with their targeted promoters. Further refinement of this protocol, such as the use of other restriction enzymes or differential ways to cut the DNA could improve sequence capture length. Reduction in costs in long read sequencing could also allow to generate bigger libraries and allow for ccMPRA to become more routinely used. To assay variants, cells from a pool of different individuals could be used for capture, or captured individually and mixed together, as was previously done using the self-transcribing active regulatory region sequencing (STARR-seq)^95,96^. Our capture approach could allow to readily generate MPRA libraries from cells from different organisms and/or under different treatment conditions. For the latter, it would be interesting to analyze conditions that affect chromatin organization.

Our ccMPRA identified 73.6% CREs that were not previously annotated by ENCODE SCREEN^14^ v3 that were enriched for known regulatory element histone modifications but at lower levels than ENCODE annotated sequences. As we are measuring significant regulatory activity in our assay and derive them from chromatin interactions with an active promoter region, this could be due to a higher proportion of false negatives in their identification in SCREEN, which relies on high signals of chromatin accessibility and histone modifications. However, some proportion could also be due to the capture of spurious interactions with closed chromatin or due to the capture of poised interactions that are not regulatory active in the cell type.

Capturing of promoter fragments with their respective cCRE allowed us to measure not only enhancer activity but also silencers. Due to substantial higher activity of the captured promoters compared to minimal promoter setups, our ccMPRA identified a greater number of silencers than enhancers. This allowed us to identify the following silencer-associated features: 1) promoters that interact with silencers exhibit greater dependence on orientation than promoters interacting with enhancers; 2) silencers tend to lie closer to their target promoters than enhancers; 3) weak promoters tend to have more enhancers and strong promoters more silencers. Analyses of both enhancers and silencers found them to be enriched for known enhancer and silencer marks respectively. In addition, analysis of TF binding found enhancers to be enriched for activating TFs and silencers for repressive ones, with both having many TFs with a dual role. It is important to note that some aforementioned activators or repressors could have a dual role that is yet to be discovered. For example, PRDM10 is predicted to also be part of a repressor complex^97^. The many dual activity TF enrichment of enhancers and silencers suggests that many distal CREs could be either enhancer or silencers based on context, for example the chromatin state of the promoter they interact with, which would explain why they are hard to separate^98^. This is consistent with previous reports showing that 22–62% of candidate silencer elements identified in one cell type can function as enhancers in at least one other cell type^50,53^.

Our ccMPRA suggests that CREs can have very different activity depending on their target promoter. Previous work that mixed different promoters with different CREs from three different mouse loci and measured their regulatory activity found similar results, whereby CREs have different activities depending on the promoter they are matched with^22^. Another study that carried out a *Drosophila* genome-wide STARR-seq with a housekeeping promoter and a developmental promoter found differences in enhancer activity due to the assayed promoter^99^. This was in line with an additional study that carried out STARR-seq to test the combined activity of 1,000 candidate enhancers and 1,000 promoter sequences in human K562 cell lines^23^. This study proposed a multiplicative model where the activity of enhancers and promoters combined determines regulatory activity and further observed that, owing to their high basal activity, housekeeping promoters are less responsive to enhancer effect, such that the same enhancer elicits a smaller relative fold-change on these promoters than on promoters of variably expressed genes^23^. These results are in line with ours, showing that differences in activity of enhancers are dependent on the target promoter. As we were also able to assay silencers, we observed that this characteristic seems to extend to silencers and interestingly some sequences can behave as enhancers with one promoter and as a silencer with another. In addition, the aforementioned multiplicative model also fits with our observation that weak promoters tend to have more enhancers while strong promoters mainly have silencers.

Our study has several limitations. Similar to other MPRAs, it tests sequences outside their genomic context. However, it allows to test sequences alongside their target promoters and tests longer sequences, overcoming issues with sequence length in MPRAs which generally test shorter sequences via oligosynthesis which are also more costly to generate. However, as sequences are not synthesized, it hinders the ability to test synthetic sequences, selected sequence variants, sequences from extinct species or ones that don’t have available cell lines or tissues. On the other end, it can test various inputs that can lead to changes in chromatin interactions and their effect on regulatory activity in cells by measuring both. Due to the complexity of these libraries and current high costs of long read sequencing, we tested only 650 promoters and also had to exclude several sequences that had a low number of barcodes. Further improvements of this technology, including reduction in long read sequencing costs could overcome these hurdles. In summary, we developed a novel MPRA technology that provides a ‘two for the price of one’ result, having both chromatin interaction information and regulatory activity, overcoming the caveats of using shorter sequences in MPRA and using a minimal promoter. We hope that further developments of this technology will lead to an increased understanding of the regulatory code and grammar and also how changes in chromatin interactions can affect regulatory activity.

## Methods

### Cell culture

HepG2 cells (ATCC, # HB-8065) were grown in Eagle’s Minimum Essential Medium (MEM) (Corning, 10-009-CV) supplemented with 10% FBS heat-inactivated (Gibco, A38400-01). Before use, HepG2 cell culture plates were coated with Collagen coating solution, using 1 ml per 10 cm^2^ culture area (Sigma-Aldrich, 125-50) for 30min at 37°C. LentiX 293T cells (Takara, 632180) were grown in Dulbecco’s Modified Eagle Medium (DMEM) containing Glutamax and pyruvate (Gibco, 10569069) and supplemented with 10% FBS heat-inactivated (Gibco, A38400-01). All cells were grown at 37°C with 5% CO2 and were regularly tested for absence of mycoplasma contamination using the LookOut® Mycoplasma PCR Detection Kit (Sigma-Aldrich, MP0035). Cell types were authenticated using the ATCC STR authentication method.

### Capture Hi-C promoter probe design

RNA-seq data for HepG2 cells was retrieved from the ENCODE project (experiment accession: ENCSR329MHM). PLAC-seq H3K4me3 data for HepG2 cells was obtained from GEO (accession: GSE161873). Promoters involved in 3D contacts in the PLAC-seq data were identified using GREAT analysis^100^ (Association rule: Basal+extension: 5,000 bp upstream, 5,000 bp downstream, 0 bp max extension, curated regulatory domains included). Promoters with more than three significant interactions in both PLAC-seq replicates were identified, yielding a set of 971 promoters. These promoters were ranked based on their average expression levels, calculated from the two RNA-seq replicates. From this ranked list, the top 499 expressed promoters were selected. Among these, 55 promoters overlapped with the top 1,000 activity promoters from a previous MPRA in HepG2^94^, serving as positive controls. Additional 50 positive controls were included, which corresponded to the top 50 active promoters from a previous MPRA^94^. 53 negative controls were added corresponding to the bottom active promoters from the previous MPRA^94^, that exhibited no significant interactions in either PLAC-seq replicates as well as 48 other negative controls that were the lowest-expressed promoters, determined from the RNA-seq data, overlapping with the bottom 1,000 active promoters of the previous MPRA and with no significant PLAC-seq interactions. A total of 650 targeted TSS were selected, comprising 444 test promoters, 105 positive controls and 101 negative controls.

### Probe Design

Probes were designed to target each selected TSS, spanning a region of 500 bp upstream of the TSS to 100 bp downstream. The RNA probes were 120 bp long and were designed by Arima Genomics with the following parameters: 3X tiling density, moderate repeat masking, GC boosting and each probe being present three times in the library. As a result, the number of probes varies depending on the targeted regions and is increased in GC rich regions for a total of 34,417 probes. The final list of probes is attached in **Supplementary Table 1**.

### Promoter Capture HiC

Promoter capture Hi-C was performed using the Capture-HiC+ kit (Arima Genomics, Custom Capture module, Tier 2) with a custom-designed promoter capture panel. HepG2 cells (10 million) were dissociated and crosslinked with 2% formaldehyde (Sigma-Aldrich, 47608) in PBS 1X for 10 minutes, followed by quenching with Stop Solution 1 (provided in the kit) for 5 minutes. The cells were then stored as dry pellets containing 1 million cells per aliquot at −80°C.

#### Hi-C

The assay was done using one aliquot (1 million cells) and following the manufacturer’s protocol with modifications tailored to the ccMPRA application. Cell lysis was performed to obtain nuclei, and DNA was digested with the restriction enzyme A1 (not A2) for 30 minutes at 37°C. After ligation, reverse crosslink, and DNA purification with 0.45X AMPure XP beads (Beckman Coulter, A63881), the DNA was fragmented using a Covaris S220 focused ultrasonicator (parameters [Cycles per Burst: 200; Duty Factor: 2%; Peak Incident Power: 140 W; Treatment Time: 17sec]). A single-sided purification was then performed with 0.45X AMPure XP beads. Two biotin-streptavidin pull-down reactions were performed per sample, each starting with 1 µg of DNA (instead of the recommended 200 ng). The pulled-down DNA was subjected to library preparation, including end repair, A-tailing, and adapter ligation with a final concentration of 0.75 µM pre-annealed custom adapters 1 and 2 (**Supplementary Table 2**). The two reactions from the same sample were pooled and split into five separate PCR amplification reactions (to avoid saturation), each using P1 and P3 primers (**Supplementary Table 2**) using the Hercule polymerase (provided in the kit) for 10 cycles. The PCR conditions were modified to extend the hybridization step to 1 minute and the elongation step to 3 minutes. The amplified library was purified using 1X AMPure XP beads.

#### Capture enrichment

All the amplified DNA was pre-cleared and submitted to the capture enrichment (∼3ug instead of the recommended 1ug) according to the protocol “Arima Custom Panel (<3Mb total region size)”. To block the custom adapter sequences during the capture step, Custom xGen Blocking Oligos from Integrated DNA Technologies were used as follows: 1uL left blocker + 1uL right blocker + 3uL of Arima’s blocker from the kit. The pull-downed DNA was split into three separate PCR amplification reactions (to avoid saturation), each using P1 and P2R primers (**Supplementary Table 2**) using the Hercule polymerase (provided in the kit) to add a 15 bp random sequence serving as barcode to each P-CRE pair for 13 cycles. The PCR conditions were modified to extend the hybridization step to 1 minute and the elongation step to 3 minutes. The amplified library was purified using 1X AMPure XP beads.

### Design of candidate regulatory sequences for the standard lentiMPRA with a minimal promoter

All cCREs between 200 and 270 bp in length were selected for oligosynthesis to be tested in a standard lentiMPRA with a minimal promoter, resulting in a total of 6,231 sequences. None of these sequences overlapped known promoter regions. To have an estimate of the baseline MPRA activity, we randomly selected 200 sequences which were shuffled using MEME suite’s fasta-shuffle-letters (v 5.5.6) with default parameters. In addition, we selected 100 positive controls and 100 negative controls of a length of 270 bp that were shown to have high or low activities respectively in a previous MPRA in HepG2 cells^94^.

### Standard lentiMPRA with minimal promoter library

The lentiMPRA library was constructed as described previously^25^. Briefly, the oligo pool was amplified via a 5-cycles PCR (primers 5BC-AG-f01 and 5BC-AG-r01, **Supplementary Table 2**) using the NEBNext® Ultra™ II Q5® Master Mix (New England Biolabs, M0544) to add the minimal promoter and spacer sequences downstream of each sequence. The amplified fragments were purified using 0.85X AMPure XP beads and subjected to an 11-cycles second-round PCR (primers 5BC-AG-f02 and 5BC-AG-r02, **Supplementary Table 2**) to add a 15 bp random sequence serving as a barcode followed by a last purification with 0.75X AMPure XP beads.

### Cloning and transformation

The pLS-*Sce*I (Addgene, 137725) plasmid was linearized with *Age*I (New England Biolabs, R3552) and *Sbf*I (New England Biolabs, R3642) digestion at 37°C for 1 hour then purified with 0.65X Ampure XP beads. The C-HiC library was cloned into the pLS-SceI by recombination using 500 ng of pLS-SceI, 400 ng of C-HiC library and NEBuilder HiFi DNA Assembly Master Mix (New England Biolabs, E2621) at 50°C for 30 min according to the manufacturer’s protocol then purified with 0.6X Ampure XP beads. The Standard minimal promoter lentiMPRA library was cloned into the pLS-SceI by recombination using 1ug of pLS-SceI, 250 ng of LentiMPRA library and NEBuilder HiFi DNA Assembly Master Mix at 50°C for 1 hour according to the manufacturer’s protocol then purified with 0.65X Ampure XP beads. The cloned libraries were electroporated into NEB 10-beta electrocompetent bacteria (New England Biolabs, C3020) (400 ng of Capture HiC library into 400 uL of bacteria and 68 ng of Standard minimal promoter lentiMPRA library in 100 uL of bacteria) using a Gemini X2 electroporator (settings [voltage: 2 kV; resistance: 200 Ohms; capacitance: 25 uF; number of pulses: 1; gap width: 1mm]). Colonies were grown overnight on carbenicillin plates and subsequently processed using a Midiprep Kit (Qiagen, 12945). Approximately 22 million colonies were collected for the C-HiC library. 1.5 million colonies were collected for the Standard minimal promoter lentiMPRA library ensuring an average of >100 barcodes were associated with each sequence.

### Barcode association sequencing

#### Standard minimal promoter lentiMPRA library

To identify the random barcodes and their corresponding sequences, the sequence-mP-barcodes fragment was amplified from 500 ng of plasmid library using primers containing flow cell adapters (P7-pLSmP-ass-gfp and P5-pLSmP-ass-i#; **Supplementary Table 2**) for 7 PCR cycles using the NEBNext® Ultra™ II Q5® Master Mix (New England Biolabs, M0544) then purified with 0.6X AMPure XP beads. The amplicons were sequenced using a Novaseq X Plus 1.5B 300 with custom primers (Read 1: pLSmP-ass-seq-R1; Read2: pLSmP-ass-seq-R2; Index1: pLSmP-ass-seq-ind1; Index2: pLSmP-rand-ind2; **Supplementary Table 2**) and with the following run settings (Read1,Index1,Index2,Read2): 146, 15, 10, 146.

#### Capture HiC library long read sequencing with Complete Genomics

To identify the random barcodes and their corresponding P-CRE pairs, P-CRE-barcodes were amplified from 2 ug of plasmid, divided in 16 PCR reactions (to avoid saturation) (primers stcLFR17c-mRNA2 and 183-TSO; **Supplementary Table 2**) using the NEBNext® Ultra™ II Q5® Master Mix (New England Biolabs, M0544) for 7 cycles. The amplicons were purified using 0.6X AMPure XP beads and then digested with *Sph*I (New England Biolabs, R3182) to linearize the contaminant plasmid template. The amplicons were size selected using the Blue Pippin system (Sage Science) with the 0.75% Dye-Free Agarose Gel Cassette and External Marker S1 (Sage Science, BLF7510) and DNA ranging from 500 bp to 3 kb (Settings: High Pass and low voltage 1-6kb) was selected to eliminate the contaminant plasmid template. The eluted DNA was purified with 1X AMPure XP beads. cLFR library construction and sequencing: 8 pg of PCR product from the amplification of Hi-C templates was further amplified in a 75 µL PCR mix containing 37.5 uL KAPA HiFi HotStart Uracil+ Ready Mix (Roche Diagnostics, Indianapolis, IN), 500 pM of each primer, 1M betaine, and 500 pM dUTP. The PCR mix was incubated for 3 minutes at 98°C followed by 15 cycles of 15 seconds at 98°C, 30 seconds at 62°C, and 5 minutes at 72°C. The PCR product was purified with a 1x volume of AMPure XP beads (Beckman Coulter, Indianapolis, IN). The purified sample was end-blocked and uracil digested in 50 µL of 1× Terminal Transferase Buffer (New England Biolabs, Ipswich, MA) containing 250 µM CoCl2, 80 units Terminal Transferase (New England Biolabs, Ipswich, MA), 200 µM ddATP, 20 units UDG (New England Biolabs, Ipswich, MA), and 40 units APE1 (New England Biolabs, Ipswich, MA). The reaction was incubated for 30 minutes at 37°C, then purified with a 1× volume of AMPure XP beads (Beckman Coulter, Indianapolis, IN). Next, 3 µM concentration of branch adapter was ligated to the uracil digested sample with 400 units T4 ligase (New England Biolabs, Ipswich, MA) in 50 µL ligase buffer containing 50 mM Tris-HCl (pH 8.3), 10 mM MgCl2, 0.05 mg/mL BSA, 1 mM ATP, and 10% PEG-8000. The reaction was incubated with 54 cycles of 30 seconds at 15°C and 30 seconds at 37°C, then purified with a 1x volume of AMPure XP beads (Beckman Coulter, Indianapolis, IN). The ligation product was amplified in a 75 uL PCR mix containing 37.5 uL KAPA HiFi HotStart Ready Mix (Roche Diagnostics, Indianapolis, IN) and 500 pM of each primer. The PCR mix was incubated for 3 minutes at 98°C followed by 7 cycles of 15 seconds at 98°C, 30 seconds at 62°C, and 3 minutes at 72°C. The PCR product was purified with a 1x volume of AMPure XP beads (Beckman Coulter, Indianapolis, IN). PCR product was denatured in a 30 µL reaction with 10 µM splint oligo, and 100 mM NaOH. Following a 5 minute incubation at room temperature, the reaction was neutralized with 3 µL of 1 M Tris-HCl buffer (pH7.0). Single strand circularization was performed with 600 units of T4 ligase in a 60 µL reaction containing 50 mM potassium acetate, 25 mM Tris-acetate (pH 7.5), 7.5 mM magnesium acetate, 2.5 mM ATP, and 0.4 mM DTT. Following ligation, circles were digested with 30 units of Exonuclease I (New England Biolabs, Ipswich, MA) and 50 units of Exonuclease III (New England Biolabs, Ipswich, MA) to remove any uncircularized product. The circles were purified with a 2.5x volume of AMPure XP beads (Beckman Coulter, Indianapolis, IN). The controlled primer extension was performed on 40 fmol of circles in a 25 µL 1x ThermoPol Buffer (New England Biolabs, Ipswich, MA) containing 400 pM primer, 6 pmol dNTP, and 1.25 units Taq DNA polymerase (New England Biolabs, Ipswich, MA), incubated 2 minutes at 95°C, 1.5 minutes at 54°C, and 10 minutes at 68°C. Next, 3 µM concentration of branch adapter was ligated to the controlled primer extension product using 400 units T4 ligase (New England Biolabs, Ipswich, MA) in 50 µL ligase buffer containing 50 mM Tris-HCl (pH 8.3), 10 mM MgCl2, 0.05 mg/mL BSA, 1 mM ATP, and 10% PEG-8000. The reaction was incubated for 1 hour at room temperature and purified with a 2x volume of AMPure XP beads (Beckman Coulter, Indianapolis, IN). The ligation product was amplified in a 60 uL PCR mix containing 30 uL KAPA HiFi HotStart Ready Mix (Roche Diagnostics, Indianapolis, IN), and 500 pM of each primer, incubated for 3 minutes at 98°C followed by 8 cycles of 15 seconds at 98°C, 30 seconds at 62°C, and 3 minutes at 72°C. The PCR product was purified with a 1× volume of AMPure XP beads (Beckman Coulter, Indianapolis, IN). For each library the PCR product was analyzed by a single lane of a flow cell on a DNBseq G400 instrument (Complete Genomics, San Jose, CA) using a paired-end 100 base CoolMPS sequencing kit (Complete Genomics, San Jose, CA). Approximately 100 Gb of data was generated per library.

#### Capture HiC library long read sequencing with Pacbio Revio

To identify the random barcodes and their corresponding P-CRE pairs, P-CRE-barcodes were amplified and purified as for Complete Genomics but with different primers for the PCR (Assoc3kb.F and Assoc3kb.R, **Supplementary Table 2**) in order to produce amplicons > 3kb. After the size selection of DNA ranging from 1 kb to 6 kb with the Blue Pippin (Settings: Pulse 2-6 kb) and purification with beads, an amplicon library was generated following standard procedures and sequencing was done with the Pacbio Revio using 3 Revio SMRT cells (∼30 million reads in total).

### Lentivirus packaging, titration and lentiMPRA infection

For lentivirus packaging, 35 million LentiX 293T cells were seeded to a 15 cm dish and directly transfected with 27.5 ug of lentiMPRA library and 22.5 ug of Virapower™ Lentiviral Packaging Mix (Invitrogen, 442050) using Lipofectamine™ 3000 transfection reagent (Invitrogen, L3000001) following the manufacturer’s protocol. The following day, the medium was refreshed. Three days post-transfection, the supernatant medium containing the lentivirus was collected, filtered through a 0.45 μm PES filter system and concentrated with 1/3 volume of Lenti-X concentrator reagent (Takara, 631232) overnight at 4°C. The following day, the supernatant was centrifuged at 1,500g for 45 min at 4°C and the pellet was resuspended in 400uL cold PBS and stored at −80°C.

For lentivirus titration, 200,000 HepG2 cells were seeded to 8 wells of a 24 wells plate in medium supplemented with polybrene 8ug/mL to increase the transduction efficiency and 0, 1, 2, 4, 8, 16, 32 or 64uL of lentivirus were added directly to each well. The following day the medium was refreshed without polybrene. The genomic DNA was extracted three days after infection, to harvest primarily integrating lentivirus, using the Wizard SV Genomic DNA Purification Kit (Promega, A2361). The multiplicity of infection was determined by qPCR, measuring the relative abundance of viral DNA (WPRE region, amplified with WPRE_F and WPRE_R primers; **Supplementary Table 2**) after subtracting the unintegrated plasmid backbone signal (BB.F and BB.R primers; **Supplementary Table 2**) and normalizing to genomic DNA (intronic region of the LIPC gene, amplified with LP34.F and LP34.R primers; **Supplementary Table 2**) for each sample, as described in Gordon et al.^25^. Reactions were conducted using SsoFast EvaGreen Supermix (Bio-Rad, 1725205) according to the manufacturer’s protocol.

For the lentiMPRA infection, the number of cells required was determined as total barcode integrations (number of candidate regulatory sequences tested x 100 barcodes x 100 integration/barcode; arbitrary set as 1,500,000,000 for the Capture-C MPRA) divided by the MOI (∼100 for HepG2 cells), using the spreadsheet in **Supplementary Table 3**. HepG2 cells were transduced in 3 technical replicates with lentivirus in a medium supplemented with polybrene 8 ug/mL. The medium was refreshed without polybrene the following day. For the Capture-C MPRA, 2 biological replicates (with 3 technical replicates each) were performed.

### lentiMPRA sequencing

The lentiMPRA sequencing library was prepared as described in Gordon et al.^25^. Briefly, three days after infection, cells were washed 3 times with PBS and DNA and RNA were extracted using the Qiagen AllPrep DNA/RNA kit (Qiagen, 80204), according to the manufacturer’s protocol. The RNA barcodes were reverse transcribed using the primer P7-pLSmP-ass16UMI-gfp and the Superscript IV Reverse Transcriptase (Invitrogen, 18090010) according to the manufacturer’s protocol. cDNA and DNA barcodes were amplified by PCR for 3 cycles (primers P7-pLSmP-ass16UMI-gfp and P5-pLSmP-5bc-i#, **Supplementary Table 2**) to add sample index and UMI then purified with 1.4X AMPure XP beads. The barcodes were amplified a second time using P5 and P7 primers for 10-15 cycles then purified with 1.2X AMPure XP beads. The LentiMPRA sequencing libraries were sequenced on an Illumina Novaseq X Plus 1.5B 100 or with Novogene using the Illumina Nextseq High-output using custom primers (Read1: pLSmP-ass-seq-ind1; Read2: pLSmP-bc-seq; Index1: pLSmP-UMI-seq; Index2: pLSmP-5bc-seq-R2; Supplementary Table 2) and with the following run settings (Read1,Index1,Index2,Read2): 15, 16, 10, 15.

### De novo assembly of Complete Genomics long fragment reads

Hi-C fragments incorporated in the lentiMPRA library were determined using multiple sequencing approaches for establishing an association of incorporated sequences and their MPRA tags, as described above. One approach used Complete Genomics’ complete long fragment read (cLFR) technology, which can sequence and assemble DNA molecules up to ∼5 kb by generating 100 bp short reads that are linked to the original fragment through barcode sequences. First, all barcodes with less than 50 reads were removed. The remaining reads were quality trimmed using BBTools (Bushnell B. – <u>sourceforge.net/projects/bbmap/</u>) bbduk.sh (parameters “qtrim=rl trimq=10”). Reads with the same barcode were grouped using sgrep (https://sgrep.sourceforge.net/) and assembled de novo using megahit^101^ (parameters “--k-min 31 --k-max 41--min-contig-len=600”), resulting in 10,361,092 sequences. The resulting sequences were used together with the PacBio HiFi (i.e. circular consensus reads) in the identification of capture Hi-C interactions described below.

### Processing of association and count sequencing files

We developed a publicly available Snakemake^102^ workflow (https://github.com/kircherlab/mpra_capture_flow) which takes association and count sequencing files as input and performs identification of capture Hi-C interactions, prepares the input files for MPRAsnakeflow (https://github.com/kircherlab/MPRAsnakeflow/) and performs activity quantification using BCalm^103^.

### Identification of capture Hi-C interactions

#### Trimming

Each association sequence is separated from its barcode by a linker sequence which we removed using cutadapt v4.4^104^. The sequences were obtained as one single sequence, either in the order 5’ adapter - sequence - linker - barcode - 3’ adapter, or in the order 3’ adapter - barcode - linker - sequence - 5’ adapter. Based upon whether the linker was found in forward or reverse orientation, the sequences were saved as forward or reverse. We used seqtk seq v1.4 (https://github.com/lh3/seqtk) to reverse all reverse sequences and barcodes to ensure the same orientation as in the MPRA construct library. We filtered all sequences for a minimum length equal to the length of the 3’ adapter minus 50. Some of the constructs erroneously contain multiple linker sequences. After removing linkers and combining forward and reverse sequences, these appear as duplicates which are removed before further processing. The 5’ adaptors attached to the sequence (**Supplementary Table 2**) were removed using bbduk.sh. The 3’ adaptors, which are attached to the barcode, were removed by extracting the barcode as the first 15 bp and discarding the remaining sequence.

#### Mapping

The sequences were cut into fragments using the religation sequence for DpnII (GATC), where we kept the GATC at the ends of both sequences. The separate fragments were then mapped to the human GRCh38 genome build (as provided by the University of California Santa Cruz (UCSC) Genome Browser^105^ as hg38) using bwa mem2 v2.2.1^106^. We removed all fragments that contained more than two different interacting sequences (long Hi-C fragments can contain multiple ligations, also known as Hi-C walks^107^). From the remaining mapped fragment sequences, we excluded all supplementary alignments, accepted hard clippings and trimmed soft clippings. All sequences that did not contain a religation site were saved in a separate file for later analysis.

#### Identifying interactions

We quantified and filtered valid interactions using the filter function in HiCUP v0.9.2^26^. HiCUP filter requires a digested genome file, which we created using hicup digester on the hg38 genome for DpnII. From the filtered interactions we calculated the capture efficiency using HiCUP capture.

#### Calling significant interactions

We used CHiCAGO^29^ to identify significant loops in the cHi-C data. To do this, we first used chicagoTools to create rmap, baitmap and so-called design files from the digested genome file using a resolution of 5kb. We merged mapped sequences from different association sequencing rounds using samtools merge. This merged file was converted to a chinput file, the input format for CHiCAGO, using chicagoTools. The chinput file was used to identify significant loops (a CHiCAGO score > 5) with CHiCAGO.

#### Grouping sequences into bins

We binned all mapped fragments that overlapped at both sides of a loop. To do so, we first added information indicating in which order two interacting fragments were sequenced using tags in the BAM alignment lines. Either CO:Z:left or CO:Z:right are added to each sequence in the merged bam file from multiple association sequencing rounds. These tags were then used to filter out only left- or only right-sided interactions using samtools view^108^. We overlapped these subsets with the digested genome using bedtools^109^ intersect with a minimum overlap fraction of 0.7. To merge the sequences into bins, we first join the left and right sides using GNU Coreutils 9.9.1 join on the original sequence IDs. Then all Hi-C fragments overlapping the same restriction fragment are binned using GNU awk 5.1. After sorting these by their coordinates using GNU Coreutils, adjacent restriction fragments containing the same Hi-C fragments are merged into larger bins using GNU awk.

### MPRA count processing

#### Preparing association input

The trimmed fastq files (see “Identifcation of capture Hi-C interactions - trimming” described above) of all Hi-C sequences and barcodes of the different association runs within the same biological replicate are merged. Next, the barcodes are concatenated to the sequence ID using the attachBCToFastQ python script from MPRAsnakeflow. All duplicate sequences are marked using BBtools^110^ dedupe.sh (parameters “minidentity=99 maxedits=10 minlengthpercent=99 minoverlappercent=99 mergenames=t”). Each marked duplicate is assigned the sequence ID of the first duplicate in the group of duplicates, and a unified fasta file is written with the new sequence ID, the barcode, and the consensus sequence of the duplicates. This fasta file is then used to create a barcode assignment file containing the corresponding sequence ID of each barcode. We used MPRAsnakeflow’s filterAssignmentTsv python file to filter barcode collisions with a minimum support of 1 sequence per barcode and a minimum fraction of 0.6 to support assignment. The deduplicated fastq file, after unification of IDs, is formatted into a fasta file to use as a design file for MPRAsnakeflow.

#### Count processing

The MPRA count tables were obtained using MPRAsnakeflow’s experiment workflow, using the design and assignment files created as described in the previous step. No minimum threshold for barcodes per associated sequence was used.

### Statistical analysis

#### Activity quantification

We calculated the log RNA/DNA (log ratios) and p-values using BCalm^103^. To this end, we merged barcode count tables of the two lentiMPRA biological replicates (see “LentiMPRA sequencing library” described above). This table was joined with the file containing the sequence to bin assignments (described as the output of the “Grouping sequences into bins” step described above) using the Python package Polars (see workflow). We filter each barcode to have an MPRA RNA count of at least one in at least three of the total six replicates (of both MPRA experiments combined). Further, we required for each bin to have at least five representative barcodes. To run BCalm on this data, it was reformatted into separate RNA and DNA files, with barcode counts per replicate as columns, and bins as rows.

#### Annotation of regions

For further analysis we annotated the paired bin regions in a Python notebook using Polars (https://github.com/kircherlab/CMPRA_figures CMPRA5_add_labels_OA.ipynb). We used GENCODE^111^ v44 GRCh38 gene model annotations to obtain transcriptional start sites (TSSs) and ENCODE SCREEN^14^ v3 for GRCh38 for known candidate Cis-Regulatory Element (cCREs) annotations. We further annotated the paired bins with the BCalm output (logFC, adjusted p-value), the distance from the target promoter, the target promoter’s gene and whether the target promoter is a target promoter, a positive control or a negative control promoter.

#### Z-score calculation

The z-score was obtained by subtracting the logFC of a promoter-cCRE interaction with the logFC of the promoter only, divided by the sample standard deviation of the difference between each cCRE interacting with the same promoter. All cCREs with a z-score higher than 2 were labeled as enhancers for that promoter, and all cCREs with a z-score less than −2 were labeled as silencers.

#### Region enrichment analysis

All enrichment analyses were performed using the R package regioneReloaded v1.8^112^. We used the following publicly available assay data, with ENCODE accession numbers: DNase-seq (ENCSR149XIL), H3K36me3 ChIP-seq (ENCSR000DUD), H3K79me2 ChIP-seq (ENCSR000AOM), H2AFZ ChIP-seq (ENCSR000AOK), H3K4me1 ChIP-seq (ENCSR000APV), H3K9ac ChIP-seq (ENCSR000AMD), H3K27ac ChIP-seq (ENCSR000AMO), H3K4me2 ChIP-seq (ENCSR000AMC), H3K9me3 ChIP-seq (ENCSR000ATD), ATAC-seq (ENCSR042AWH), H3K27me3 ChIP-seq (ENCSR000DUE), H3K4me3 ChIP-seq (ENCSR575RRX), H4K20me1 ChIP-seq (ENCSR000AMQ), Hi-C (ENCSR194SRI). We further obtained all available TF ChIP-seq data in HepG2 cells from ReMap 2022^47^.

#### Comparison to regular MPRA

The regular MPRA with minimal promoter was processed using MPRAsnakeflow to get count tables and BCalm to get logFC activity values. We were able to recover 5,963 out of 6,231 CRE sequences from the test group, 199 out of 200 from the shuffled controls, 87 out of 100 from the negative controls and 60 out of 100 from the positive controls. A normalized activity score was calculated by subtracting the median activity of the shuffled control group and dividing the result by the standard deviation of the shuffled control group. This normalized score was subtracted from the ccMPRA z-score to get a promoter specificity index (PSI) score. We classified cCREs as having a low, medium or high PSI score by setting the lower 5th and upper 95th percentile of all PSI scores as thresholds. For easier interpretation, we filtered the high PSI scores to have a ccMPRA z-score of at least 1, and the low PSI scores to have a ccMPRA z-score of at most −1.

### TFBS motif enrichment

#### Motif search and scanning

We looked for both known and *de novo* transcription factor binding site (TFBS) motifs in our set of silencers and enhancers using HOMER v2^56^ findMotifsGenome.pl and compared to a random genomic background composed by HOMER. We used the HOCOMOCO v13^113^ clustered non-redundant motif database for evaluating known motifs. We additionally scanned all CRE and promoters separately for known motif presence using findMotifsGenome.pl with the “-find” option. We removed all matches with a score lower than 6.

#### Logistic regression single motifs

The results from the motif scanning were used to create a binary feature input set to perform a lasso logistic regression on (https://github.com/kircherlab/CMPRA_figureslasso_regression.ipynb). Each TFBS motif occurring in a CRE is a feature, and each motif in a promoter is a feature. If an interaction has the same motif in CRE and promoter part, it would have a 1 in the CRE column for that motif and a 1 in the promoter column for that motif. The sequence class is 1 if the interaction contains an enhancer and −1 if it contains a silencer. We used both chromosomes 17 and 6 as holdout chromosomes to use as a test set, because this combination resulted in the best class balance with 888 positive and 914 negative sequences in the training set and 273 positive and 243 negative sequences in the test set. We used scikit-learn v1.6.1 to train a logistic regression model with elastic net regularization with an inverse regularization strength C=0.009 and an L1 ratio of 0.1, which we obtained using cross validation. The trained model achieved an AUROC score of 0.65, and retained 233 of the original 1,159 features. From these selected features, 135 features attributed more to the positive class (enhancers) and 112 to the negative class (silencers). For both enhancers and silencers, there are slightly more promoter motifs than CRE motifs. Interestingly, enhancer-promoter interactions share more motifs between the enhancer and promoter part (11% of motifs are shared) than do silencer-promoter interactions (5% of motifs are shared).

#### Logistic regression motif combinations

From the single motif logistic regression selected features, we took the 50 most important motifs of both classes. We created a new input feature set with all possible combinations between promoter and CRE motifs. We again trained a logistic regression model with elastic net regularization and cross validation, resulting in an inverse regularization strength C=0.04 and and L1 ratio of 0.8. The trained model had an AUROC score of 0.63 on the test set, and retained 50 of the original 2,680 input features. Each of these 50 features represents a co-occurrence of two motifs, one in the CRE and on in the promoter part of an interaction, that contributes to the readout MPRA activity.

## Acknowledgements

We thank Elphege Nora (UCSF) for inspiring this study, the ENCODE Consortium and the ENCODE production laboratories for generating the datasets used here, Allyson Whittaker and Anthony Schmitt from Arima Genomics for their assistance and advice on capture Hi-C, and Max Schubach and Kilian Salomon for their input and advice for the data analysis. This work was supported in part by the National Human Genome Research Institute (NHGRI) grant numbers 1R21HG010065 (Y.S., N.A.), UM1HG009408 (J.S., N.A.) and UM1HG011966 (J.S., M.K., N.A.) and the UCSF Sandler Program for Breakthrough Biomedical Research, which is partially funded by the Sandler Foundation (Y.S., N.A.). C.A. was supported in part by EMBO (ALTF 585-2021), the Bakar Aging Research Institute postdoctoral fellowship and the Bettencourt Schueller foundation.

## Author contributions

C.A, Y.S. and N.A. conceived the study. C.A., F.I., X.C. devised the methodology; C.A., X.C., K.A., and L.M. carried out experimental work; X.Z., R.D., B.A.P. carried out sequencing via Complete Genomics, C.A. and P.K. analyzed results; P.K. wrote custom analysis software and processed all data; J.S., Y.S., M.K., N.A. supervised the project and acquired funding. C.A, P.K., M.K. and N.A. wrote, reviewed and edited the paper with contributions from all authors.

## Data availability

Processed analysis files of this Capture-C MPRA experiment can be found at https://doi.org/10.5281/zenodo.15511615.

Hi-C data in HepG2 was retrieved from ENCODE (accession ENCSR194SRI).

ATAC-seq data in HepG2 cells was retrieved from ENCODE (accession ENCSR042AWH).

All ChIP-seq data in HepG2 cells was retrieved from ENCODE and ReMap2022^47^.

## Code availability

The source code is available at GitHub (https://github.com/kircherlab/mpra_capture_flow/ and https://github.com/kircherlab/CMPRA_figures/).

---

**Extended Data Fig. 1:**
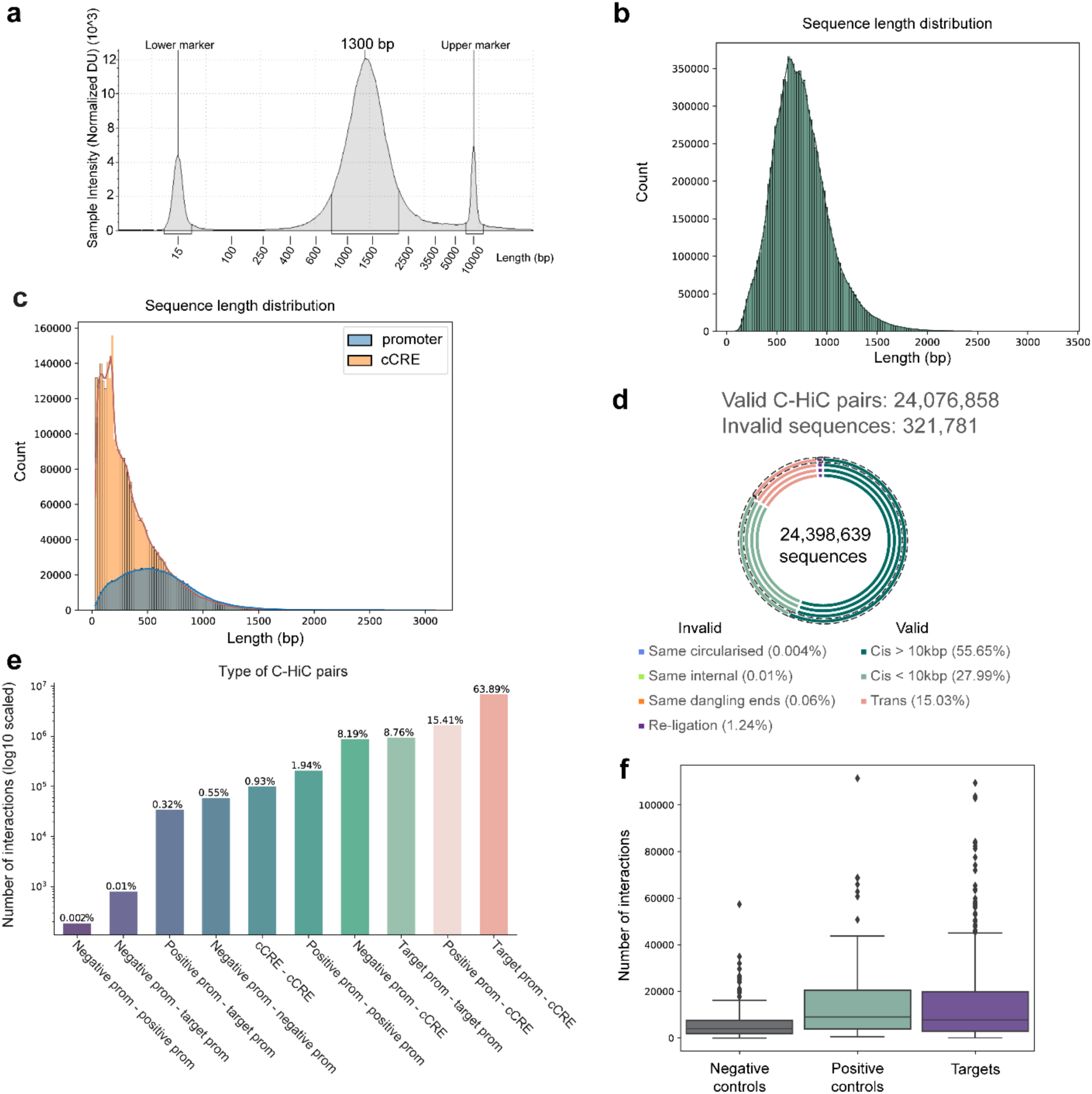
Validation of promoter C-HiC. **a**, Tapestation profile representing the length distribution of the DNA after sonication used for the C-HiC. **b**, Distribution of the total length of the C-HiC pairs (in bp). **c**, Distribution of the length of each region of the C-HiC pairs: the captured promoter versus the cCRE regions. **d**, Proportion of valid C-HiC pairs (containing *trans* or *cis* interacting regions) and invalid sequences determined with the HiCUP tool v0.9.2^26^. The four circles represent the results obtained from the four long read sequencing runs (3 Pacbio Revio and 1 Complete Genomics (outer dashed circle)). The average percentage of the four sequencing runs is indicated for each category. **e**, Breakdown of the type of interactions found in the captured Hi-C pairs (positive prom = positive control promoter; negative prom = negative control promoter; target prom = target promoter of interest). **f**, Number of interactions per promoter for each promoter category. The median is shown and the lines on either side of the box plot represent the 1.5 interquartile range (IQR).

**Extended Data Fig. 2:**
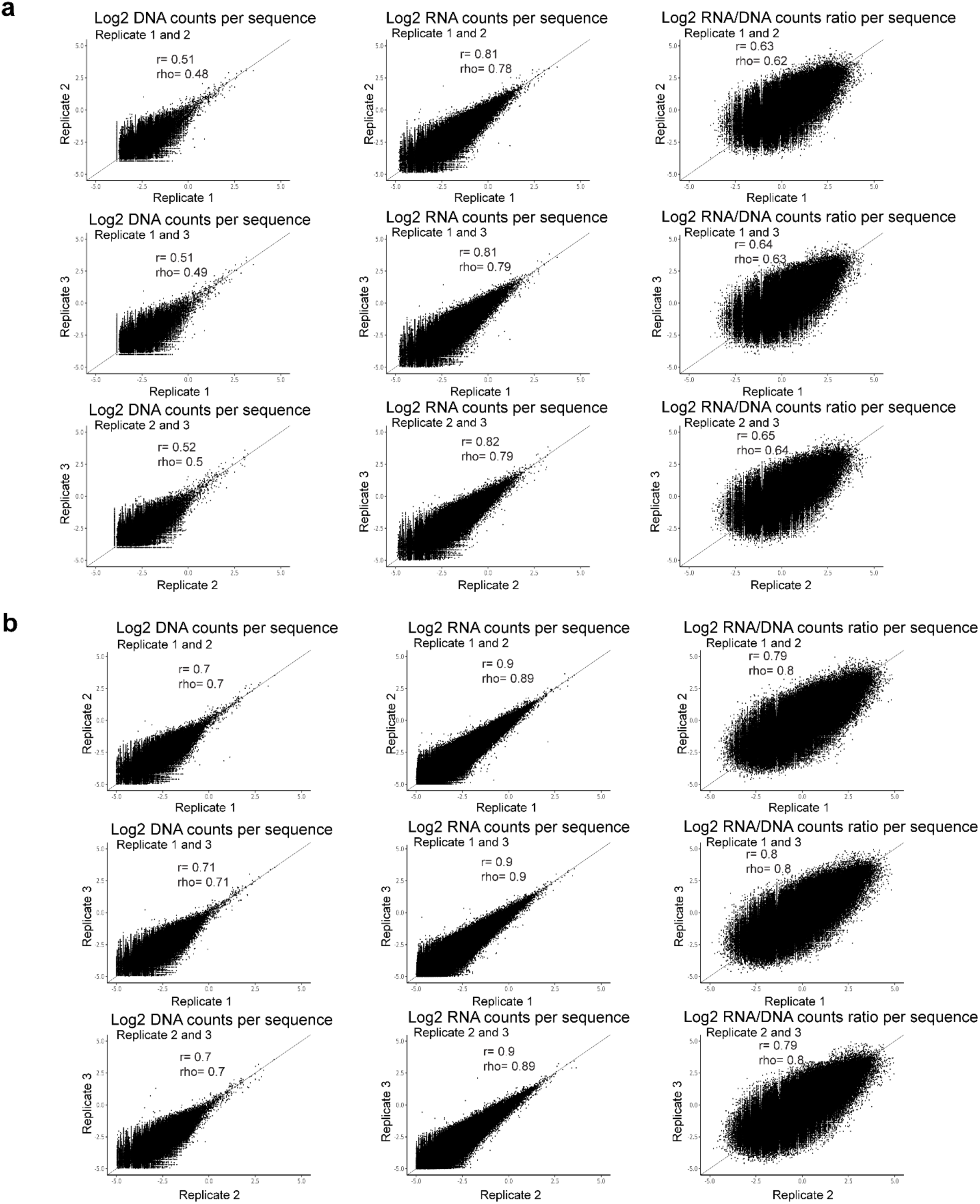
Reproducibility among ccMPRA replicates. **a-b**, Scatter plots displaying the relationship between observed barcode DNA counts, RNA counts and RNA/DNA ratios per C-HiC sequence for all pairwise comparisons among replicates of two independent MPRA sets (performed using two different lentiviral libraries) (**a** and **b**). The Pearson (r) and Spearman (rho) correlation coefficients are also provided.

**Extended Data Fig. 3:**
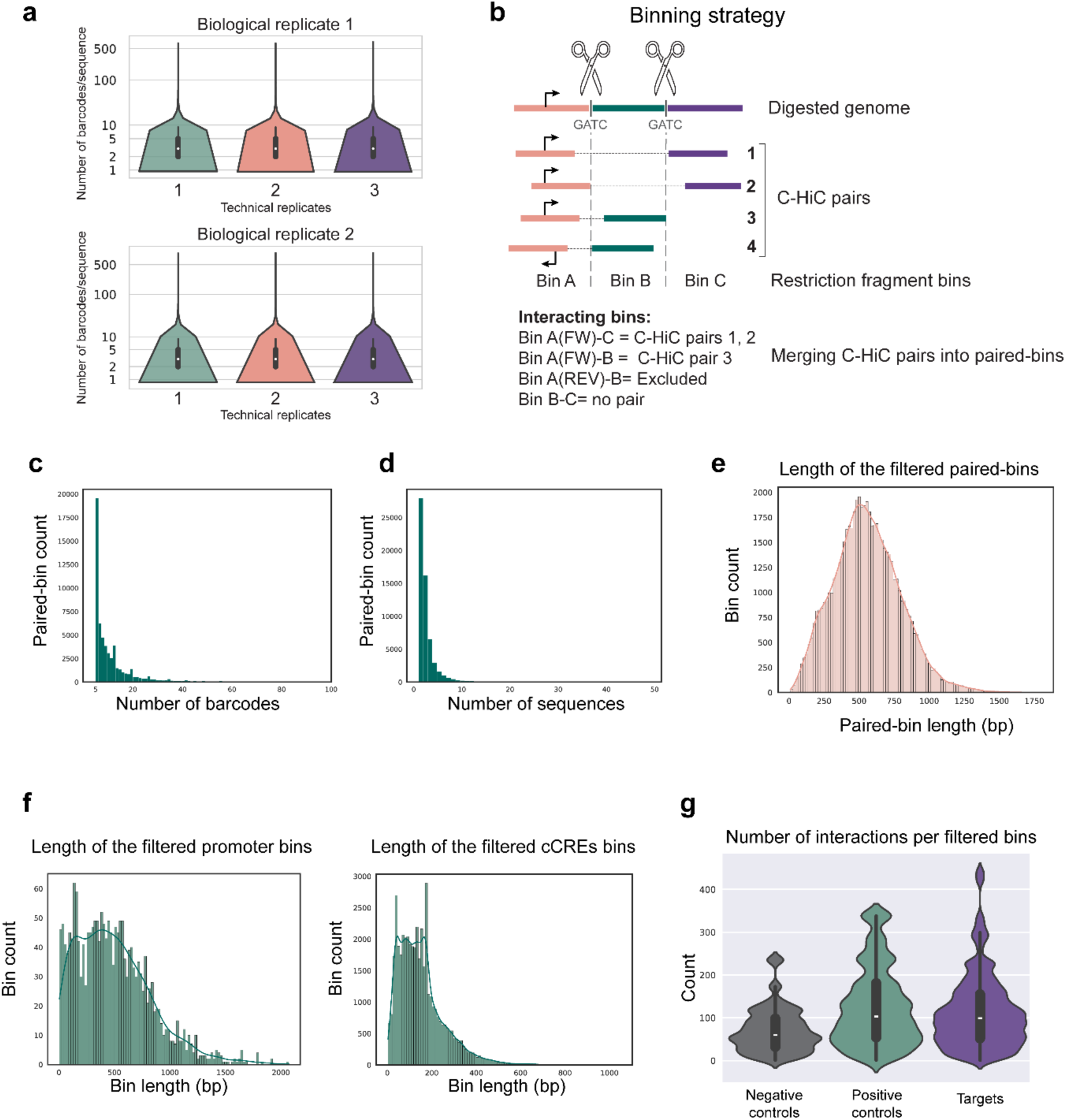
Binning and filtering of the C-HiC pairs. **a**, Barcodes per sequence containing a bait promoter detected in the MPRA for two MPRA biological replicates, each containing three technical replicates. The median is shown with a white dot on a black bar representing the 1.5 interquartile range (IQR) in each violin plot. **b**, The genome is digested *in silico* using the restriction site ‘GATC to produce restriction fragments that were used as bins. The C-HiC pairs mapping to the same pairs of bins are merged into a paired-bin under the condition that the promoter is equally oriented. Thus, promoters in the forward (FW) orientation are in different bins than promoters in the reverse (REV) orientation. **c**, Distribution of the number of barcodes per paired-bin containing a bait promoter after filtering out the bins associated to <5 barcodes. **d**, Distribution of the number of DNA sequences per filtered paired-bin containing a bait promoter. **e**, Length distribution of filtered paired-bins containing a bait promoter in base pairs (bp). **f**, Distribution of the length (in bp) of the filtered bait promoter (left) and cCREs (right) bins. **g**, Violin plots showing the Absolute number of interactions per promoter bin for each promoter category. The median is shown as a white dot on a black bar representing the 1.5 interquartile range (IQR) of each violin plot. For an independent t-test, positive vs negative: t(17215) statistic= 43.42300, p-value< 0.0001; target vs negative: t(51589) statistic= 38.64034, p-value<0.0001; target vs positive: t(56202) statistic= - 11.70100, p-value= 1.37046e-31.

**Extended Data Fig. 4:**
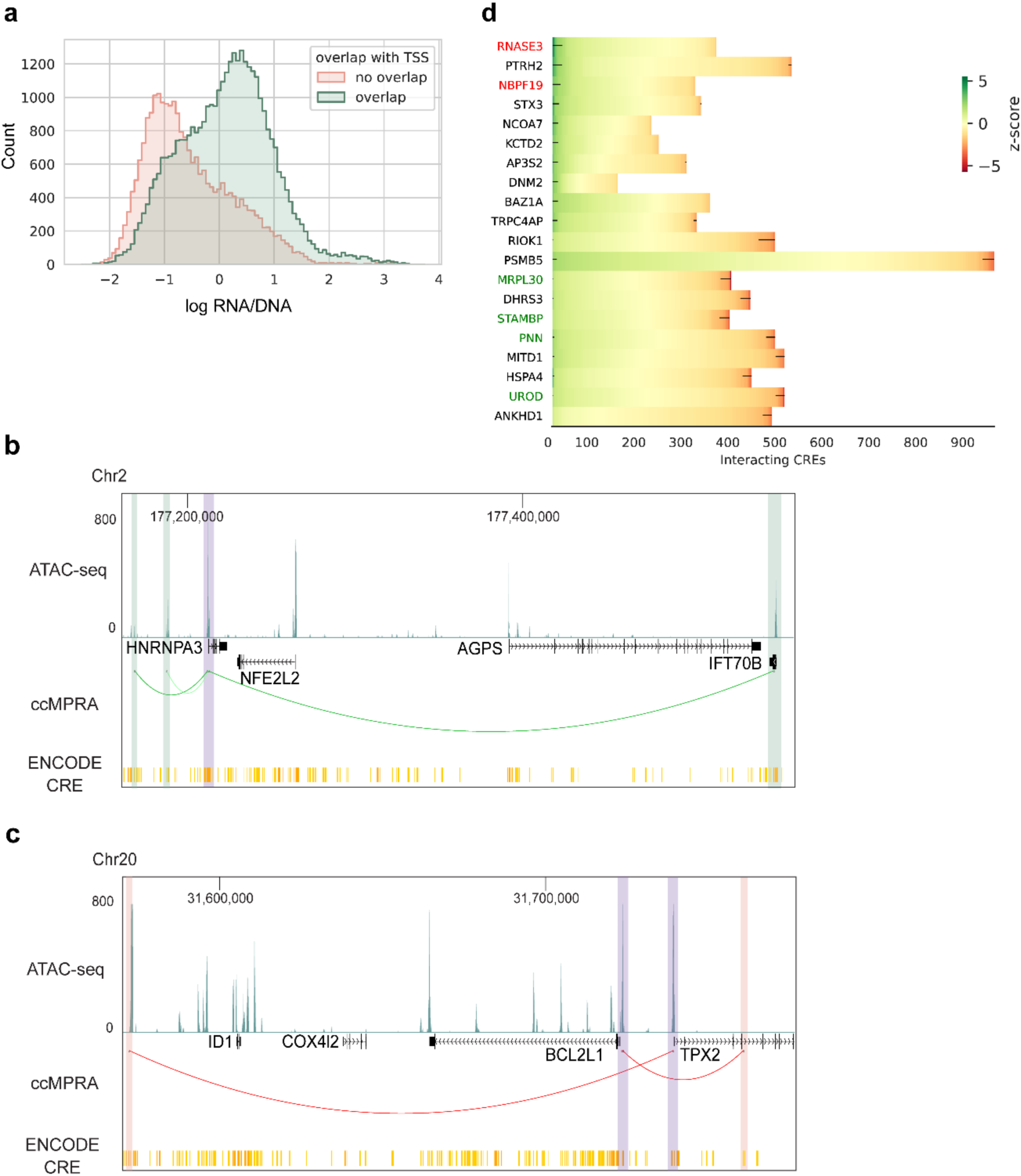
lentiMPRA regulatory activity of C-HiC sequences. **a**, Plot displaying the distribution of the lentiMPRA activity of C-HiC pairs determined as the log2(RNA/DNA) ratios and split between those overlapping with a transcriptional start site (TSS, green) or those that are not (red). **b-c**, UCSC Genome Browser^38^ screenshot showing examples of an enhancer identified by ccMPRA for *HNRNPA3* (**b**) or a silencer for *BCL2L1* and *TPX2* (**c**). HepG2 ATAC-seq and CREs from ENCODE^31^ SCREEN v3 are shown alongside our ccMPRA signal (green and red arcs represent an enhancing or silencing interaction, respectively). Purple shade indicates target promoters and the green/red shade depicts the enhancer or silencer regions. **d**, Heatmap showing the distribution of *cis* regulatory element (CRE) bins across a subset of promoters, along with their associated regulatory activity (z-score). Each row represents a promoter with the gene name in the y-axis, and each column corresponds to a CRE bin. Black dots/lines along the rows indicate CRE bins classified as significant enhancers or silencers, when their z-score is >2 or <-2 respectively. Promoters in red, green or black font indicate negative controls, positive controls and target promoters, respectively.

**Extended Data Fig. 5:**
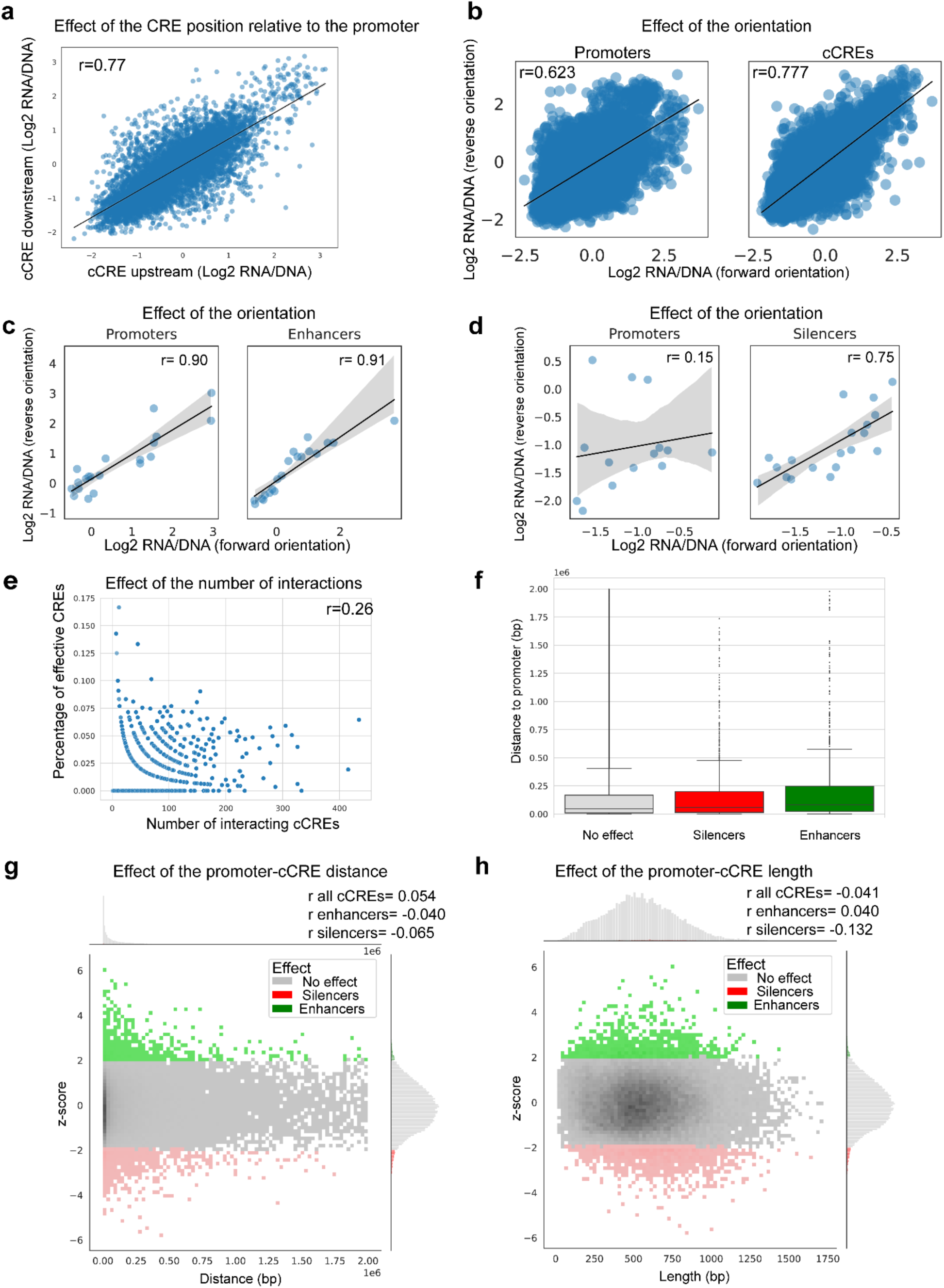
Architectural determinants of CRE activity. **a**, Correlation of lentiMPRA activity between upstream and downstream positions of cCREs relative to the promoter in the lentiMPRA library. The Pearson correlation is shown in the upper left corner. **b**, Correlation of lentiMPRA activity between forward and reverse orientations of promoters (left panel) or cCREs (right panel) in the lentiMPRA library. The Pearson correlations are shown in the upper left corners. **c**, Correlation of lentiMPRA activity between forward and reverse orientations of the promoters interacting with enhancers, where enhancers are averaged across both orientations (left panel) or the correlation between forward and reverse orientations of their enhancers, with promoter orientation averaged across both orientations (right panel) in the lentiMPRA library. The Pearson correlations are indicated in the upper right corners. **d**, Correlation of lentiMPRA activity between forward and reverse orientations of the promoters interacting with silencers, where silencers are averaged across both orientations (left panel) or the correlation between forward and reverse orientations of their silencers, with promoter orientation averaged across both orientations (right panel). **e**, Correlation between the number of interacting regions (cCRES) per promoter and the percentage of active CREs (regions that possess a regulatory activity). The Pearson correlation is shown in the upper right corner. **f**, Distribution of the endogenous distance between cCREs and their target promoter whether they have no regulatory effect (grey), are silencers (red) or enhancers (green). For an independent t-test, enhancer vs no effect: t(55594) statistic= 6.11239, p-value= 9.87962e-10; enhancer vs silencer: t(2135) statistic= 4.20220, p-value= 2.75211e-05; silencer vs no effect: t(55959) statistic= 0.70160, p-value= 0.48292. The median is shown and the lines on either side of the box plot represent the 1.5 interquartile range (IQR). **g-h**, Correlation between cCREs z-scores and their genomic distance to their target promoter (**g**) or length (**h**) for sequences that have no regulatory effect (grey), enhancers (green) or silencers (red). Histograms are shown outside of the plot area. Pearson correlations for the highlighted groups are shown in the upper right corner.

**Extended Data Fig. 6:**
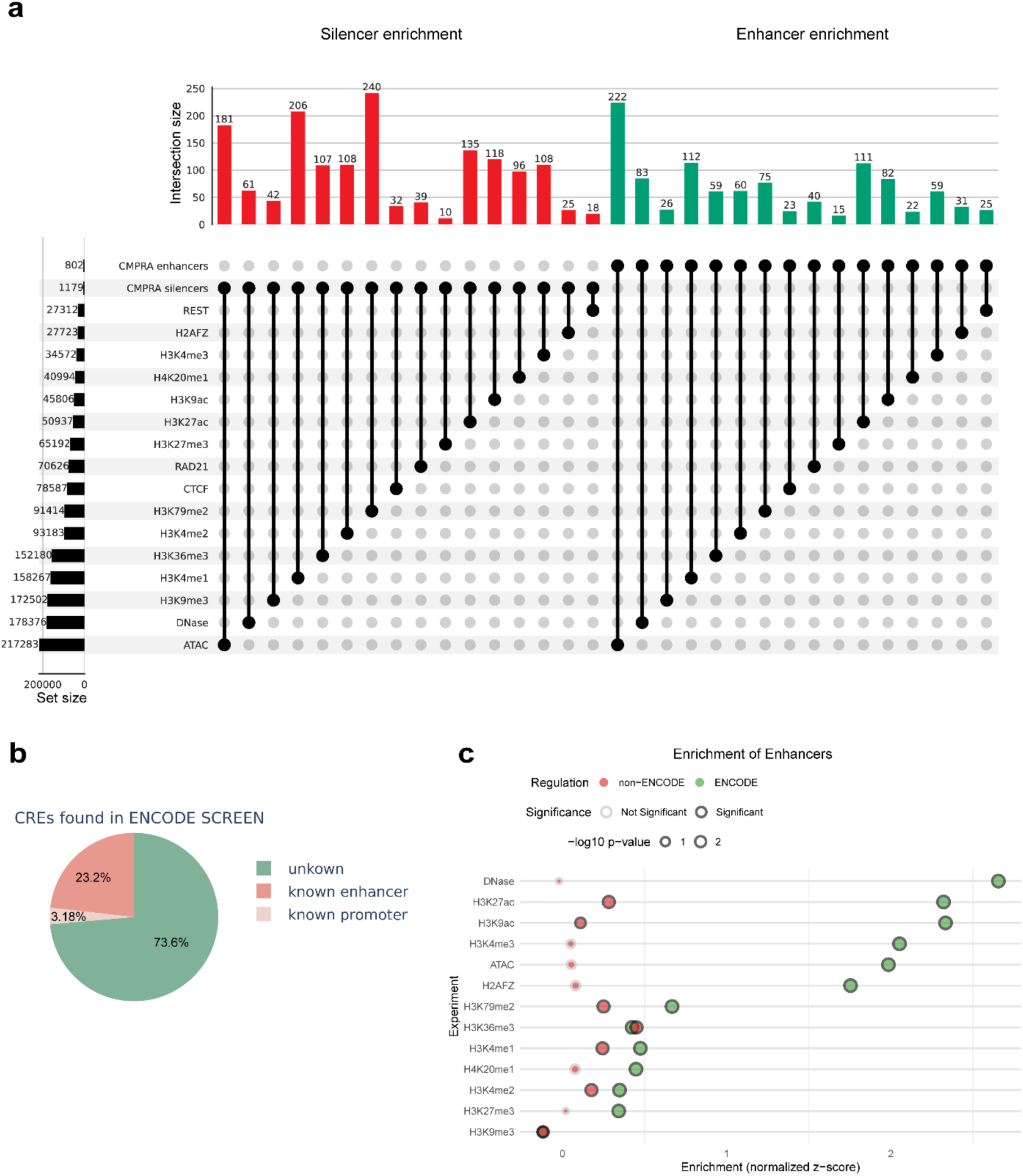
Features of ccMPRA CREs. **a**, Upset plot showing the number of silencers (left, red) or enhancers (right, green) identified by ccMPRA overlapping with various chromatin modifications or transcription factors (ChIP-seq data from ENCODE^31^ and ReMap 2022^53^). **b**, Percentage of the CREs identified by ccMPRA, overlapping with ENCODE^31^ SCREEN v3 annotated CREs. **c**, ATAC-seq, DNAse hypersensitive sites and ChIP-seq (histone modifications) signal enrichment from ENCODE^31^ (HepG2 cells) is shown across enhancers identified by ccMPRA that overlap with SCREEN annotated CREs (green) or are not annotated in SCREEN (red). Circle size represents -log10 p-value of permutation tests.

**Extended Data Fig. 7:**
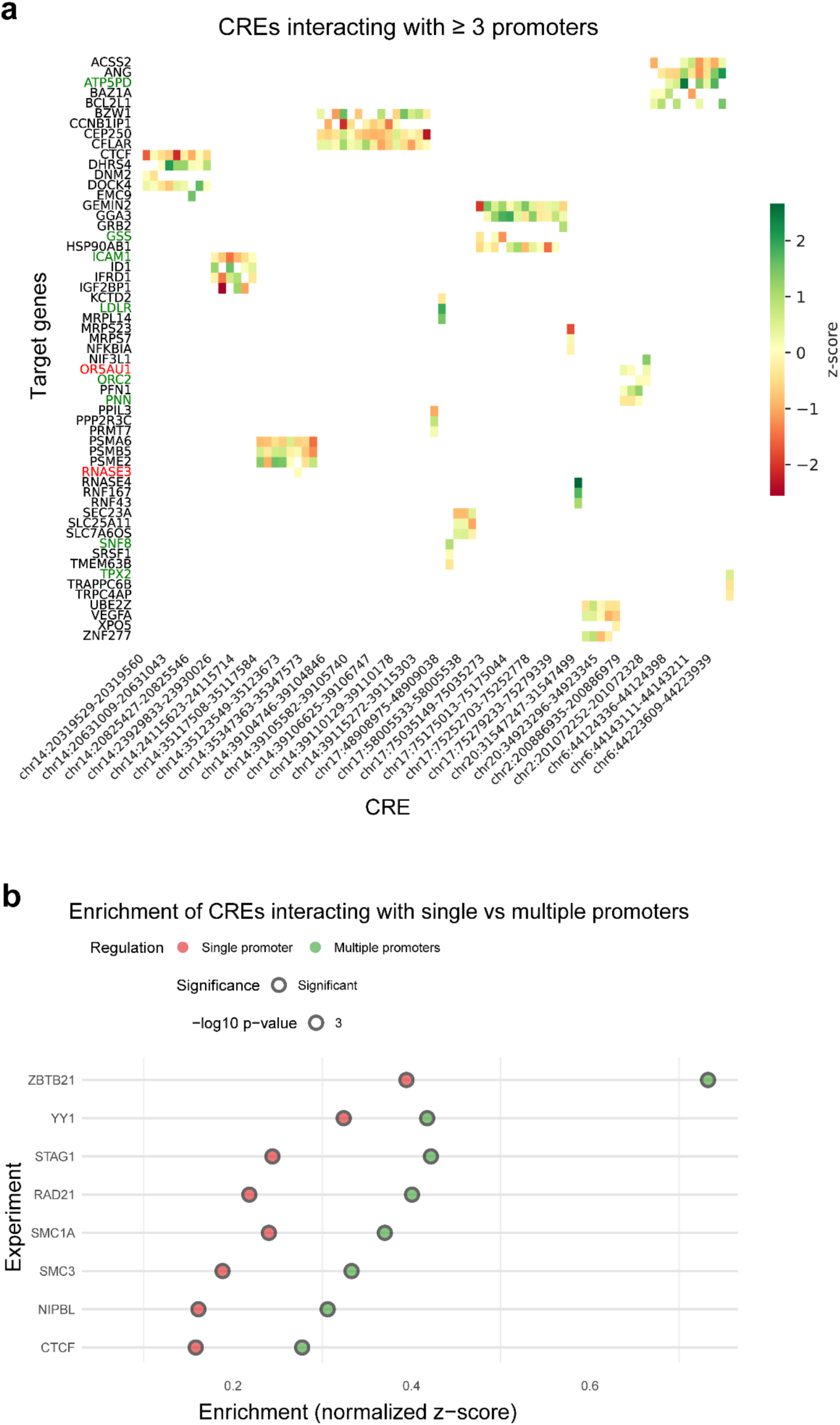
Promoters affect CRE activity. a, Heatmap showing the regulatory activity (z-score) of CREs interacting with three or more promoters, plotted separately for each promoter they contact. b, Enrichment of ChIP-seq signal analyzed from ReMap 2022^53^ (HepG2 cells) for various transcription factors across CREs identified by ccMPRA, whether they interact with a single promoter (red) or with multiple promoters (green). Circle size represents -log10 p-value of permutation tests.

**Extended Data Fig. 8:**
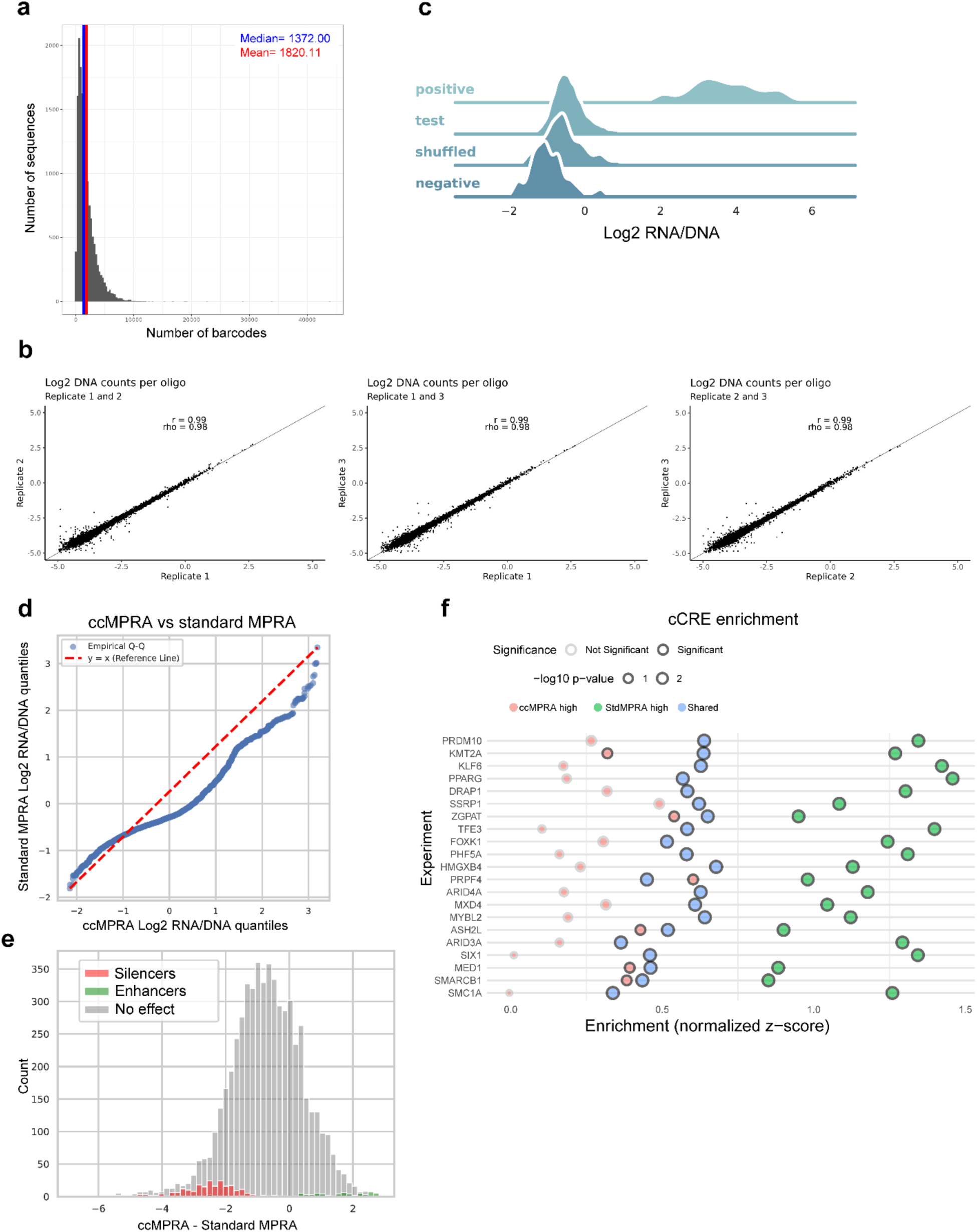
Comparison of ccMPRA to a standard minimal promoter MPRA. **a**, Barcodes per sequence obtained in the standard MPRA using a minimal promoter. **b**, Scatter plots displaying the relationship between observed barcode DNA counts, RNA counts and RNA/DNA ratios per MPRA sequence for all pairwise comparisons among three technical replicates. The Pearson (r) and Spearman (rho) correlation coefficients are also provided. **c**, Distribution of log2(RNA/DNA) ratios for positive and negative control sequences, as well as ccMPRA-identified cCREs (test) and shuffled sequences. **d**, QQplot showing the quantile values of the ccMPRA activity plotted against the quantile values of the standard MPRA activity for all tested sequences. **e**, Distribution of sequences based on their differential activity scores between the two assays (ccMPRA z-score – stdMPRA z-score). Sequences identified as enhancers or silencers by ccMPRA are highlighted in green and red, respectively. **f**, Enrichment of ChIP-seq signal analyzed from ReMap 2022^53^ (HepG2 cells) for various transcription factors across CREs identified by ccMPRA, whether they have a higher detected activity in the ccMPRA (ccMPRA high, red), in the standard MPRA (StdMPRA high, green) or have a similar activity between the two assays (Shared, blue). Circle size represents -log10 p-value of permutation tests.

