## Supplementary Figure 1 for "Capture-C MPRA: A high-throughput method to simultaneously characterize promoter interactions and regulatory activity"

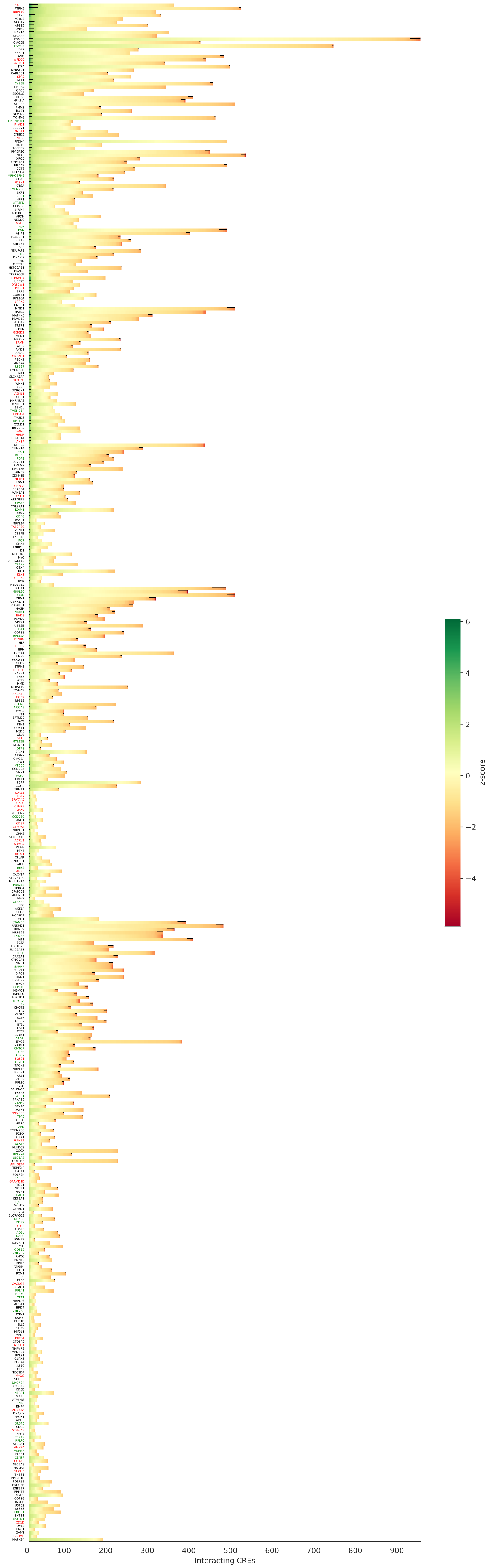


**Supplementary Figure 1: cCRE regulatory activity per promoter.** Heatmap showing the distribution of cis regulatory element (CRE) bins across all 650 promoters, along with their associated regulatory activity (z-score). Each row represents a promoter with the gene name in the y-axis, and each column corresponds to a CRE bin. Black dots/lines indicate CRE bins classified as significant enhancers or silencers, when their z-score is >2 or <-2 respectively. Promoters in red, green or black font indicate negative controls, positive controls and target promoters respectively.
