## Supplementary Table 2 for "Capture-C MPRA: A high-throughput method to simultaneously characterize promoter interactions and regulatory activity"

### Supplementary Table 2: Primer and adapter sequences

| | Sequence |
|---|---|
| **Adapter 1** | CCCGCTCTAGACCTGCAGGCGTACACGGTAC*T |
| **Adapter 2** | /5Phos/GTACCGTGTACGGCAAAGTGAACACATCGCT |
| **Primer P1** | CTCACTCAGCCTGCATTTCTGCCAGGGCCCGCTCTAGACCTGCAGGCGTACACGGTAC*T |
| **Primer P3** | CTTAGCTTTCGCTTAGCGATGTGTTCACTTTGCCGTACACGGTAC*T |
| **IDT Left Blocker** | CTCACTCAGCCTGCATTTCTGCCAGGGCCCGCTCTAGACCTGCAGGCGTACACGGTACT/3block/ |
| **IDT Right Blocker** | CTTAGCTTTCGCTTAGCGATGTGTTCACTTTGCCGTACACGGTACT/3block/ |
| **Primer P2R** | TGAACAGCTCCTCGCCCTTGCTCACCATGGTGGCGACCGGTNNNNNNNNNNNNNNNCTTAGCTTTCGCTTAGCGATGTGTTC |
| **5BC-AG-f01** | CTCACTCAGCCTGCATTTCTGCCAGGGCCCGCTCTAGACCTGCAGGAGGACCGGATCAACT |
| **5BC-AG-r01** | GCTTTCGCTTAGCGATGTGTTCACTTTGCACAGTACCGGATTGCCAAGCTGGAAGTCGAGCTTCCATTATATACCCTCTAGTGTCGGTTCACGCAATG |
| **5BC-AG-f02** | CTCACTCAGCCTGCATTTCTG |
| **5BC-AG-r02** | TGAACAGCTCCTCGCCCTTGCTCACCATGGTGGCGACCGGTNNNNNNNNNNNNNNNCTTAGCTTTCGCTTAGCGATGTGTTC |
| **P7-pLSmp-ass-gfp** | CAAGCAGAAGACGGCATACGAGATGCTCCTCGCCCTTGCTCACCATG |
| **P5-pLSmP-ass-i#** | AATGATACGGCGACCACCGAGATCTACAC[10nt index]CAGCCTGCATTTCTGCCAGGG |
| **pLSmP-ass-seq-R1** | GGCCCGCTCTAGACCTGCAGGAGGACCGGATCAACT |
| **pLSmP-ass-seq-R2** | CATTATATACCCTCTAGTGTCGGTTCACGCAATG |
| **pLSmP-ass-seq-ind1** | GCAAAGTGAACACATCGCTAAGCGAAAGCTAAG |
| **pLSmP-rand-ind2** | TCTAGAGCGGGCCCTGGCAGAAATGCAGGCTG |
| **stcLFR17c-mRNA2** | TGTGAGCCAAGGAGTTGNNNNNNNNNNNNNNNTTGTCTTCCTAAGACCGCTTGGCCAGCCTGCATTTCTGCCAGGG |
| **183-TS0** | GAGACGTTCTCGACTCAGCAGAGCTCCTCGCCCTTGCTCACCATG |
| **Assoc3kb.F** | GGCAAAGAGAAGAGTGGTGC |
| **Assoc3kb.R** | AGCCAAGGAAAGGACGATGA |
| **LP34.F** | TCCTCCGGAGTTATTCTTGGCA |
| **LP34.R** | CCCCCCATCTGATCTGTTTCAC |
| **WPRE.F** | TACGCTGCTTTAATGCCTTTG |
| **WPRE.R** | GGGCCACAACTCCTCATAAAG |
| **BB.F** | TGCCGCATAGTTAAGCCAGTA |
| **BB.R** | TCAAGCCTTGCCTTGTTGTAG |
| **P7-pLSmp-ass16UMI-gfp** | CAAGCAGAAGACGGCATACGAGATNNNNNNNNNNNNNNNNGCTCCTCGCCCTTGCTCACCATG |
| **P5-pLSmP-5bc-i#** | AATGATACGGCGACCACCGAGATCTACAC[10nt index]GCAAAGTGAACACATCGCTAAGCGAAAGC |
| **P7** | CAAGCAGAAGACGGCATACGAGAT |
| **P5** | AATGATACGGCGACCACCGAGATCTACAC |
| **pLSmP-bc-seq** | GCTCCTCGCCCTTGCTCACCATGGTGGCGACCGGT |
| **pLSmP-UMI-seq** | ACCGGTCGCCACCATGGTGAGCAAGGGCGAGGAGC |
| **pLSmP-5bc-seq-R2** | CTTAGCTTTCGCTTAGCGATGTGTTCACTTTGC |
